# A Community Challenge for Pancancer Drug Mechanism of Action Inference from Perturbational Profile Data

**DOI:** 10.1101/2020.12.21.423514

**Authors:** Eugene F. Douglass, Robert J Allaway, Bence Szalai, Wenyu Wang, Tingzhong Tian, Adrià Fernández-Torras, Ron Realubit, Charles Karan, Shuyu Zheng, Alberto Pessia, Ziaurrehman Tanoli, Mohieddin Jafari, Fangping Wan, Shuya Li, Yuanpeng Xiong, Miquel Duran-Frigola, Martino Bertoni, Pau Badia-i-Mompel, Lídia Mateo, Oriol Guitart-Pla, Verena Chung, DREAM CTD-squared Pancancer Drug Activity Challenge Consortium, Jing Tang, Jianyang Zeng, Patrick Aloy, Julio Saez-Rodriguez, Justin Guinney, Daniela S. Gerhard, Andrea Califano

## Abstract

The Columbia Cancer Target Discovery and Development (CTD^2^) Center has developed PANACEA (PANcancer Analysis of Chemical Entity Activity), a collection of dose-response curves and perturbational profiles for 400 clinical oncology drugs in cell lines selected to optimally represent 19 cancer subtypes. This resource, developed to study tumor-specific drug mechanism of action, was instrumental in hosting a DREAM Challenge to assess computational models for *de novo* drug polypharmacology prediction. Dose-response and perturbational profiles for 32 kinase inhibitors were provided to 21 participating teams, who did not know the identity or nature of the compounds, and they were asked to predict high-affinity binding among ~1,300 possible protein targets. Best performing methods leveraged both gene expression profile similarity analysis, and deep-learning methodologies trained on individual datasets. This study lays the foundation for future integrative analyses of pharmacogenomic data, reconciliation of polypharmacology effects in different tumor contexts, and insights into network-based assessment of context-specific drug mechanism of action.

## INTRODUCTION

Non-canonical drug targets are known to contribute to clinical toxicity (off-target effects). More recent work, however, suggests that off-targets may also drive clinical efficacy (Dar et al., 2012; Lin et al., 2019). Systematic, *de novo* elucidation of compound mechanism of action (MoA), including polypharmacology, is thus emerging as a critical, yet still highly elusive problem in clinical oncology. Availability of new methodologies for the comprehensive assessment of on- and off-target drug binding could help discriminate between targets driving efficacy or toxicity, and those producing non-relevant clinical effects (Hopkins, 2008).

Traditionally, the molecular targets of a drug that comprise its MoA have been defined by detailed thermodynamic (binding strength) and crystallographic (binding structure) characterization of a drug’s interaction with individual proteins (Anderson, 2003). This approach is quite effective as it directly facilitates structure-based drug design. Unfortunately, such a “one-drug/one-target” paradigm is often insufficient to mechanistically elucidate clinical phenotypes induced by even classical drugs, such as aspirin (Bedard et al., 2020; Proschak et al., 2019). As a result, there is an increasing need to systematically characterize drugs in terms of their polypharmacology, defined as their binding affinities across a comprehensive protein landscape (Milletti and Vulpetti, 2010). A complementary and even more complex issue is the assessment of key secondary effectors, downstream of a drug’s high-affinity binding targets, which may be highly context-specific and may thus determine the ultimate pharmacological activity of a compound in specific tissues.

An increasing number of efforts have emerged to leverage large-scale perturbational profiles—i.e., mRNA profiles of cell lines and tissues before and after perturbation with a small compound—to predict both high-affinity binding targets and context-specific effectors (Bansal et al., 2014; Iorio et al., 2010; Shen et al., 2017; Woo et al., 2015). The key assumption behind the use of perturbational profiles for this purpose is that differential gene expression is controlled by transcription factors and co-factors that represent the key downstream effectors of a compound’s high affinity binding targets (**Figure 1A**) (Alvarez et al., 2016, 2018). For example, the drug lapatinib inhibits EGFR (target) which induces gene expression changes via downstream transcription factors, including MYC and E2F family proteins (effectors) (Blumer and Johnson, 1994; Kolch et al., 2015). As a result, drug-induced differential expression of MYC and E2F transcriptional targets may help distinguish EGFR inhibitors from inhibitors with a different downstream effector repertoire (**Figure 1A**). By extension, compounds targeting the same proteins should induce similar transcriptional signatures and, vice-versa, the transcriptional signature of a compound should provide insights into its mechanism of action.

**Figure 1.**
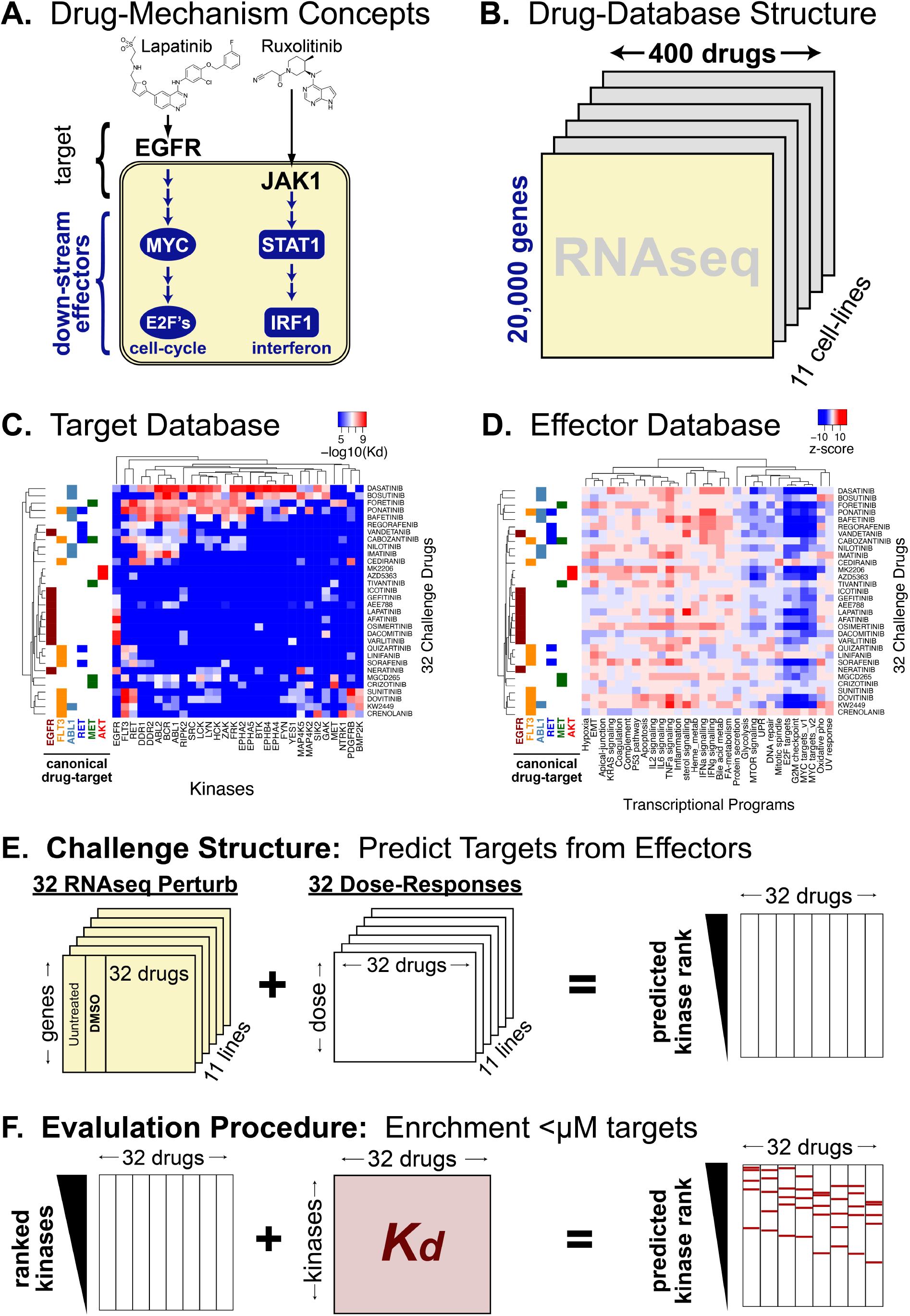
Underlying data and structure of poly-pharmacology community challenge. **(A)** Drug mechanism can be divided into direct binding targets and downstream effectors **(B)** the PanACEA database given transcriptional profiles of cell-lines perturbed by clinical oncology drugs **(C)** Kinome-binding profiles of 32 kinase inhibitors **(D)** Transcriptional Hallmark programs induced by 32 kinase inhibitors **(E) Challenge Structure:** participants are given perturbed RNAseq and dose response data and asked to predict protein-targets. **(F) Challenge Evaluation:** participant predictions are evaluated based on the enrichment of < μM binders within each drug-target prediction vector.

Availability of a compound-matched, tissue-specific dose response curve (DRC) further improves compound targets assessment. First, it allows perturbational profile generation at high, yet sub-lethal concentrations, thus preventing emergence of cell-mediated responses, such as apoptosis or cellular stress that would confound the true mechanism of action. Second, availability of differential cell killing in multiple molecularly-distinct tissues further informs on compound activity based on availability of specific proteins and complexes (Schenone et al., 2013).

Protein kinases represent one of the most thoroughly studied drug target classes. The rationale is twofold. First, protein kinases comprise some of the most frequently mutated oncogenes; as a result, their inhibitors have provided critical support for the oncogene addiction hypothesis. Second, ATP-competitive pull-down assays enable effective and systematic binding affinity measurements across comprehensive protein kinase repertoires. The most comprehensive such evaluation to date, the Kinome-Binding Repertoire (KBR), measured the affinity of 230 clinically-relevant kinase inhibitors across 255 kinases (Klaeger et al., 2017). While restricted to this protein class, this highly informative and robust dataset is well-suited to benchmarking methods aimed at predicting drug polypharmacology, thus providing much needed systematic and objective criteria for the evaluation of systems pharmacology approaches.

To leverage the KBR in assessing the research community’s ability to predict kinase inhibitors’ mechanism of action from drug perturbation profiles, we designed a DREAM Challenge (Costello et al., 2014; Saez-Rodriguez et al., 2016) based on PANACEA (Pan-cancer Analysis of Chemical Entity Activity)—a newly-developed, large-scale resource comprising genome wide RNASeq profiles and matched DRCs of multiple cell lines, following perturbation with hundreds of clinically relevant compounds. This extends and complements previous computational and systems pharmacology DREAM challenges by shifting the question from drug sensitivity to drug MoA. The PANACEA data used in this challenge includes matched dose-response curves and perturbational RNASeq profiles, representative of approximately 400 clinical oncology drugs—including FDA approved and late-stage (phase 2 and 3) compounds—on 11 cell lines, in replicate (**Figure 1B**). From these, we selected a subset of 32 kinase inhibitors that were also represented in the KBR (**Figure 1C,D**).

Challenge participants were provided with perturbational profiles and DRCs (**Figure 1E**) and were asked to predict high-affinity binding targets for the 32 drugs by developing and training machine learning algorithms on these data, on data from public databases such as (Barretina et al., 2012; Iorio et al., 2016; Subramanian et al., 2017), as well as by leveraging insights and models developed in previous DREAM Challenges (Bansal et al., 2014; Cichonska et al., 2020; Costello et al., 2014; Menden et al., 2019). To make the challenge realistic, participants were not aware that compounds had been selected from the KBR collection and that they were kinase inhibitors. The Challenge was run from December 2019 to February 2020, and led to the development and assessment of state-of-the-art approaches for inferring drug MoA from perturbational profiles, described herein.

## Results

### Challenge Requirements and Data

In the CTD^2^ Pancancer Drug Activity DREAM Challenge, participants were asked to use DREAM-provided as well as existing pharmacogenomic datasets—including cell line-matched dose-response curves (DRCs) and gene expression profiles of drug-naïve and drug-perturbed cells (perturbational profiles)—to predict compound binding proteins (high-affinity targets) of 32 anonymized drugs (**Figure 1A**). More specifically, the DREAM-provided dataset comprised 704 DCRs and matched perturbational profiles of these 32 drugs in 11 cell-lines representing molecularly distinct tumor subtypes, in replicate (**Figure 1E**). The drugs were selected so that they had molecular profile in PANACEA and experimental characterization of their high affinity binding targets in the Kinome-Binding Resource (KBR) (Klaeger et al., 2017).

Participants were encouraged to combine these data with additional, publicly available resources to infer the high-affinity binding targets of the 32 drugs from a repertoire of ~1,300 potential drug-targets. These were defined as the union of all Drug-Bank reported targets and the 255 kinases profiled in the KBR. Drug names were obfuscated, to prevent potential trivial training of the algorithm on the KBR data (**Figure 1F**), and participants were not aware that the KBR data would be used as a gold standard for performance assessment.

Consistent with past DREAM studies, the challenge included a *leaderboard round* followed by a final *validation round* (Saez-Rodriguez et al., 2016). During the former, teams were allowed to submit up to five predictions for the 32 compounds, which were scored and posted to a public leaderboard. The purpose of this round was to enable experimentation and conceptual flexibility in model development by providing rapid feedback on the accuracy of the model, while also encouraging competition among participants. A limit of 5 submissions was chosen to allow model refinement without compromising the statistical independence of the training and testing model, thus reducing the potential for over-fitting. In the final validation round, participants were asked to submit their final model’s predictions with accompanying source code, thus allowing objective validation of their methodology. Model performance was evaluated according to each team’s ability to prioritize *bona fide* targets of the 32 drugs, with the latter defined as having a dissociation constant *K_d_* < 1μM in the KBR, according to two complementary metrics, which were summarized by two sub-challenges:

***Sub-challenge 1* (*SC1*)** was designed to assess the ability of each submitted prediction to identify high-affinity binding targets (*K_d_* < 1μM) of each of the 32 compounds, among the top 10 highest scoring predicted targets. The rationale for selecting the top 10 targets was to represent the number of predictions that could be realistically validated using experimental assays. For each submitted drug prediction, a p-value was calculated by filtering the prediction list to consider only targets in the KBR, and comparing the number of *bona fide* targets (*K_d_* < 1μM in the KBR) in the top 10 predicted targets to a null model generated from all possible targets, and similarly filtered to consider only targets in the KBR. A final integrated score was computed by averaging the −log_2_(*p*-value) for each drug across all 32 drugs.

***Sub-challenge 2* (*SC2*)** was designed to assess the ability of each submitted prediction to accurately rank all the unknown *bona fide* targets (*K_d_* < 1μM), by computing their enrichment—and associated p-value—within the ranked list of predicted targets. The rationale for this second metric was to provide a more comprehensive and fine-grained comparison of the different methodologies (**Figure 1F**). Similar to SC1, a final integrated score was computed by averaging the −log_2_(*p*-value) for each drug across all 32 drugs.

### Challenge Results

During the leaderboard phase (2 months), 21 teams contributed 86 prediction matrices of which 39 (45%) showed an average mean p-value of < 0.01 via both SC1 and SC2 (**Figure S1A-B**). Interestingly, SC1 & SC2 scores revealed distinct distribution-profiles: on average, most predictions were statistically significantly enriched on the top-10-target metric (SC1), but not on the entire list enrichment (SC2) (Figure S1A-B)

Consistent with previous DREAM challenges, we established a performance ranking score for both sub-challenges by performing a bootstrap analysis of each team’s final submission by calculating a Bayes factor relative to the bootstrapped best submission for each sub-challenge (see Methods). **Figures S1C** and **2D** summarize the results of this analysis, with each box showing a team’s bootstrapped scores and the color of the box indicating the Bayes factor relative to the top performer. A Bayes factor of 3 or less indicates that models are statistically indistinguishable from the top-ranked submission. Using this criteria, *Team Atom* and *Team netphar* were confirmed as the top-performers in SC1 and SC2, respectively (**Figure S1C-D**), while team SBNB was a close second in SC2 (Bayes factor 3-5). A description of team Atom, Netphar and SBNB algorithms is provided in the STAR methods section.

To better understand the models and the difference in their performance, we examined sub-challenge scores on an individual drug basis (**Figure S1E-F**). Two clusters emerged, which separated teams based on whether they had used additional training datasets to train their algorithms. (**Table 1**). In general, overall performance was positively correlated with the number of additional databases utilized in the analysis, accounting for 27% of the variance in SC1 and a remarkable 82% of the variance in SC2.

**Table 1.** Number of Additional Datasets Used by Participants for Training & Algorithm Class

|  |  |  |  |  |  |  |  |
| --- | --- | --- | --- | --- | --- | --- | --- |
| TEAM | SC1 | SC2 | # drug-AUC datasets<sup>a</sup> | # drug-mRNA datasets<sup>b</sup> | # drug-target datasets<sup>c</sup> | Total Training datasets | Algorithm Class |
| **Netphar** | 12.6 | 70.9 | 6 | 1 | 4 | **11** | **similarity** |
| **SBNB** | 11.7 | 59.2 | 6 | 3 | 2 | **11** | **similarity** |
| **Xielab** | 13.8 | 50.3 | 6 | 2 | 1 | **9** | **similarity** |
| **Atom** | 17.4 | 49.3 | - | 2 | 4 | **6** | **NN** |
| **DMIS_PDA** | 13.8 | 35.2 | - | 2 | 1 | **3** | **NN** |
| **Theragen** | 15.1 | 17.3 | - | 2 | 1 | **3** | **similarity** |
| **Signal** | 6.3 | 6.1 | - | 1 | 2 | **4** | **regression** |
| **TeamAxolotl** | 6.2 | 1.1 | - | - | 2 | **3** | **NN** |
| **AMbeRland** | 3.3 | 1.1 | - | - | - | **0** | **unsup.** |
| **SenthamizhaV** | 7.4 | 0.9 | - | - | - | **0** | **unsup.** |

<sup>a</sup> drug-sensitivity (AUC) databases include: NCI60 (Shoemaker, 2006), GDSC (Iorio et al., 2016), CTRP (Seashore-Ludlow et al., 2015), gCSI (Haverty et al., 2016), CCLE (Barretina et al., 2012), and other manually curated data
<sup>b</sup> drug-mRNA-perturbation databases include: L1000-drugs, L1000-shRNA (Subramanian et al., 2017) and CREEDS (Wang et al., 2016)
<sup>c</sup> drug-target datasets include: DrugBank (Wishart et al., 2018), ChEMBL (Mendez et al., 2019), KEGG (Kanehisa et al., 2019), MATADOR (Günther et al., 2008)

### Contribution of Training Data Sources to Model Performances

Both SC2 winning teams, Netphar and SBNB, employed multiple highly curated datasets for training their algorithm. Netphar relied on the multi-database resources DrugComb (cytotoxicity) (Zagidullin et al., 2019) and DrugTargetCommons (drug targets) (Tang et al., 2018); SBNB relied on the multi-modality ChemicalChecker database (Duran-Frigola et al., 2020). Figure 2 provides a high-level, conceptual summary of the types of datasources included in these meta-databases, organized by data type and source.

**Figure 2.**
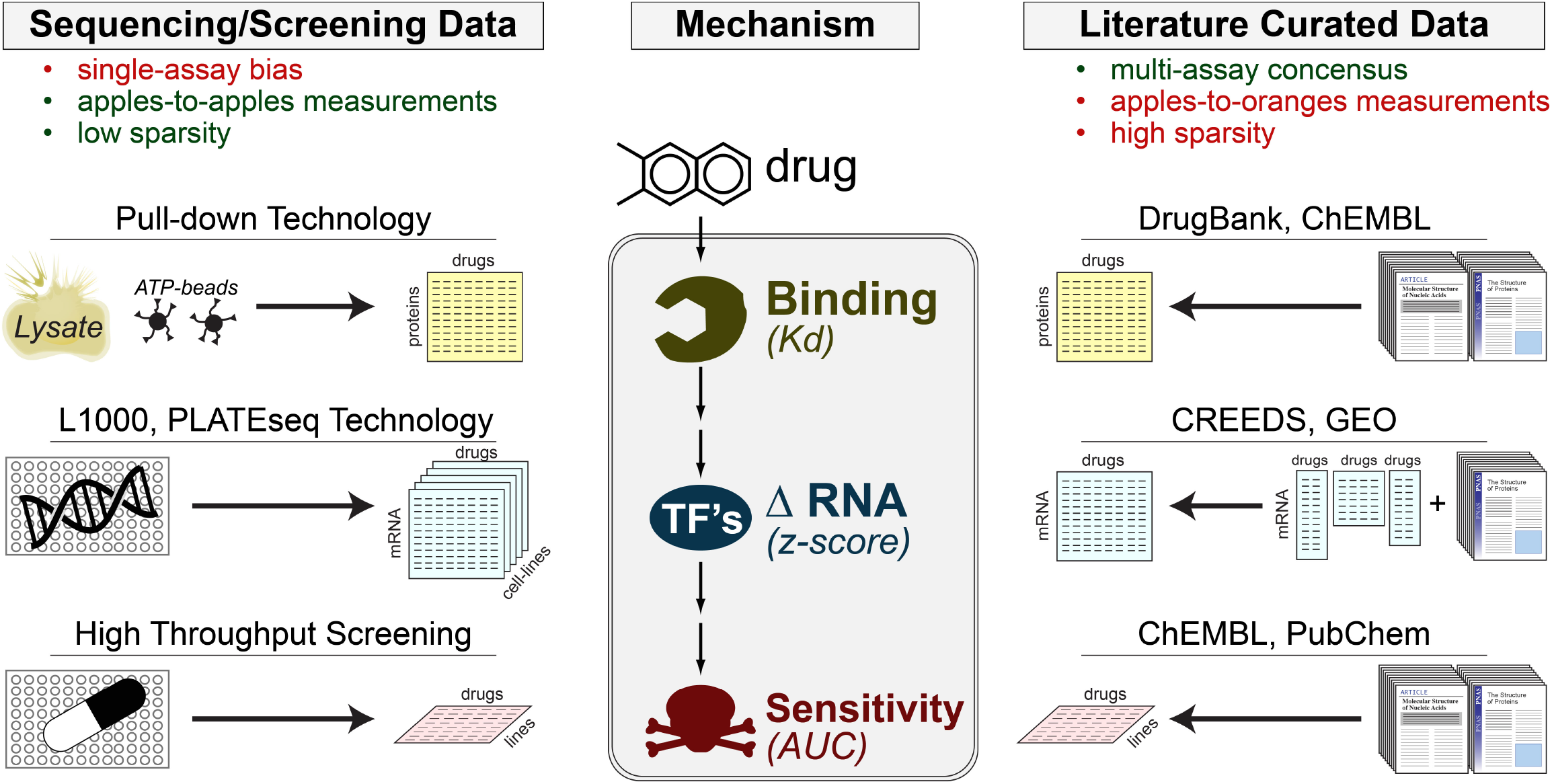
The Universe of Training Data used in this Challenge. Drug-perturbation datasets can be divided into two major categories: technology-based and literature each with distinct limitations.

Overall, the datasets used to train the algorithms could be divided into two main categories: experimental screening-based and literature curation-based (**Figure 2**). Screening approaches have the advantages of providing measurements that are quantitative, directly comparable, and systematic (i.e. low sparsity). However, they may suffer from technological platform bias. Literature-curation has the advantage of reflecting a multi-laboratory consensus but suffers from the disparate, *ad hoc* nature of the measurements and from lack of systematic assessment (high sparsity) (**Figure 2**). Team performance was further stratified based on whether they relied on (1) drug-target databases, (2) drug-perturbational databases and/or (3) cytotoxicity databases. As further discussed below, drug-target and perturbational databases provided the greatest accuracy boost across all drugs. In contrast, use of cytotoxicity databases in the training process improved performance only for a few specific drugs.

Critically, all teams chose to use literature-based datasets for identifying candidate drug targets. This is an important detail because while methods were trained on literature-based “drug-target” definitions, they were eventually evaluated based on objective, high-accuracy ATP-competitive assays (**Figure 1F**). To better understand the overlap between literature-based and ATP-based drug targets we evaluated the overlap between DrugBank and KBR targets (**Figure 3**). Specifically, we measured the number of DrugBank-reported protein kinase targets that were recovered across a range of affinity thresholds from 1 nM to 10 μM in the KBR (**Figure 3A**). Encouragingly, almost 80% of them were identified in the KBR using a *K_d_* < 1μM threshold (**Figure 3A**), consistent with a common “rule-of-thumb” for drug-lead development (Anderson, 2003). As a result, this threshold was selected to identify *bona fide*, high-affinity binding targets in the KBR for method’s performance assessment.

**Figure 3.**
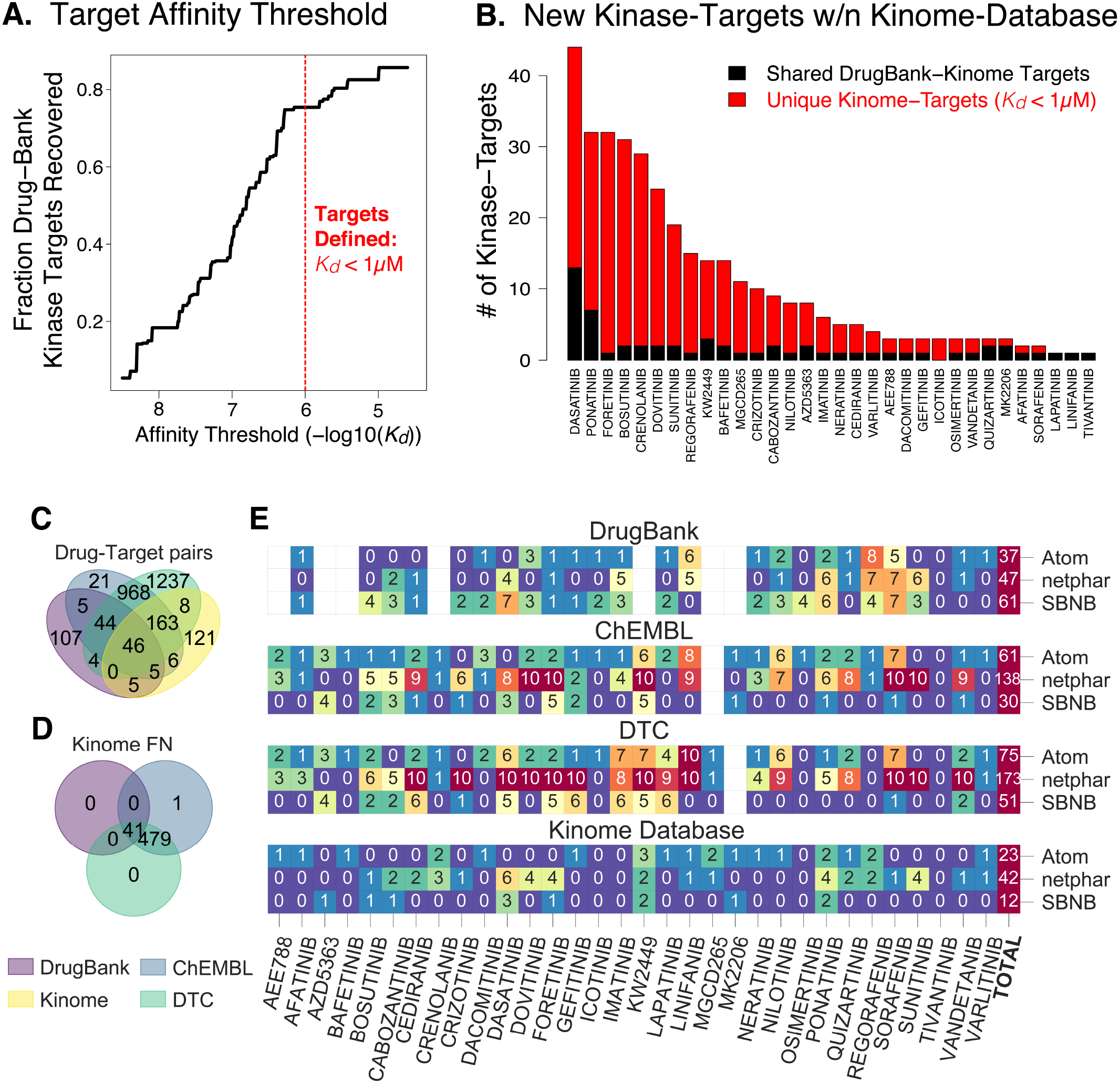
Comparison of DrugBank and Kinome Drug-Target Definitions. **A.** An affinity threshold of 1 μM within the kinome database successfully recovered almost 80% of the kinase-targets within DrugBank **B.** The Kinome-defined drug-targets appear to reveal a large number of new drug-targets (in red) in addition to the canonical drug-targets (in black) **C.** Drug-target pairs overlap across four drugtarget universes **D.** Drug-target pairs not detected in the kinome database used for PanACEA evaluation **E.** Number of successful top-10 predictions for each drug and team across the different drug target universes.

Interestingly, while a 1 μM threshold identified the majority of DrugBank kinase targets, it also revealed the presence of a significant number of new targets not reported in DrugBank (**Figure 3B**). Overall, this shows that, while DrugBank is mostly recapitulated by the KBR, the reverse is not true, suggesting that DrugBank may not contain all of the high-affinity targets of a drug. A key question raised by this comparison is whether the winning methods’ performance may have been driven entirely by canonical DrugBank Targets. To address this question, we evaluated the ratio between the scores of the top three winning teams when either DrugBank or KBR targets were used as *bona fide* high-affinity targets of the 32 drugs used in the challenge (**Figure S3**). While the scores based on DrugBank targets were consistently higher (Netphar: **3:2**, SBNB: **4:2**, Atom: **1.7:1.4**) all showed positive enrichment within the prediction vector (**Figure S3**). This result implies that literature-curated drug-targets can be successfully used to bootstrap the polypharmacology analysis of otherwise uncharacterized drugs, thus further supporting the value of these resources.

In addition to DrugBank, two additional drug target databases—ChEMBL (Mendez et al., 2019) and DrugTargetCommons (Tang et al., 2018)—were used by the top performer teams. Plotting the overlap of all drug-target pairs across all four drug-target databases, only 121 targets (34%) were found to be unique to the KBR (**Figure 3C**). Taken together, these databases provided up to 2,386 additional drugtarget interactions, of which 520 (21%) were evaluated in the KBR but were found to have affinities > 1μM, suggesting that they are false positive drug-target interactions (**Figure 3D**).

Interestingly, when comparing the overlap of the top 10 targets predicted by the winning teams in each database, the observed differences strongly reflect the training datasets used by each team (**Figure 3E**). For instance, as one would expect, SBNB and Netphar results were biased towards DrugBank and DrugTargetCommons targets, respectively.

### Kinase Groups have Distinct Transcriptional Programs

The drivers of model performance by examining prediction accuracy for individual kinase inhibitor groups, as defined in (Manning et al., 2002) (Figure 4A-B) was explored next. Significant heterogeneity in methods performance across individual drugs was observed, suggesting that differences in modeling strategies (see next section) may be leveraged to predict different drug classes. For instance, all winning methods performed better on the tyrosine-kinase inhibitors group than on any other kinase group (Figure 4C).

**Figure 4.**
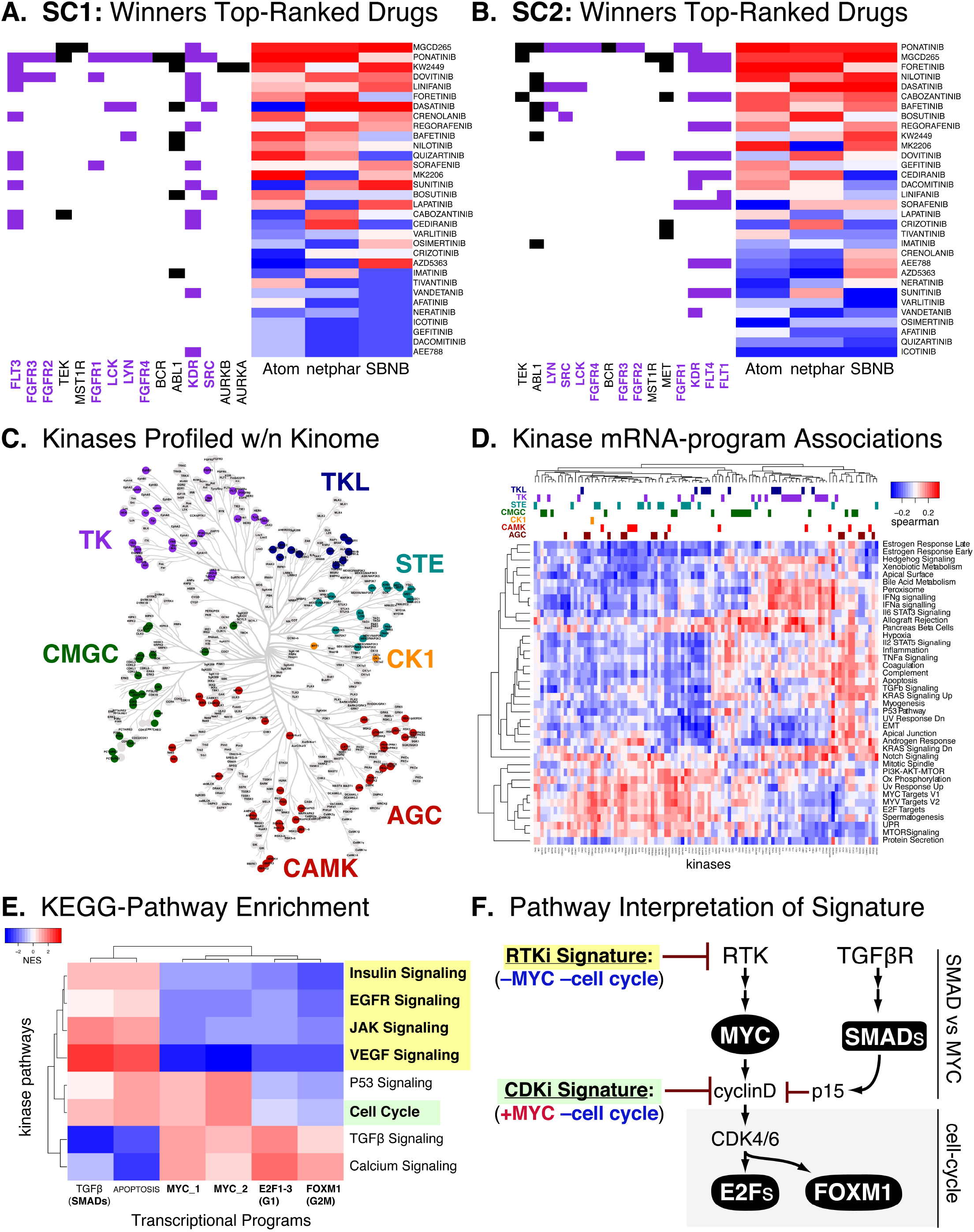
Different Kinase Pathways show distinct mRNA-signatures when inhibited. Across all models tyrosine kinase (TK) targeting drug performed the best **C.** Distribution of Kinases profiled across the Human Kinome annotated by kinase group **D.** Correlation of kinase-binding data with transcriptional-program. **E-F.** KEGG-pathway transformation of kinase-space from C revealed pathwayspecific transcriptional signatures

We thus hypothesized that specific kinase groups and families may be associated with distinct transcriptional programs. To evaluate the general relationships between kinase-targets and mRNA programs, we assessed the correlation of the KBR-reported *K_d_* with drug-induced pathway expression values (generated from the PANACEA drug perturbation database) across 84 drugs that overlap in both databases (**Figure 4D**). This correlation matrix is plotted with phylogenetic tree based kinase-groups annotated on top bars. Examining the protein kinase mRNA-program matrix, tyrosine kinases (dark purple) formed an obvious cluster (**Figure 4D columns**), showing the strongest association with proliferation programs (**Figure 4D rows, bottom cluster**): E2F targets, MYC targets, G2M checkpoint, oxidative phosphorylation, mTOR-signaling.

To better understand the nature of the biological pathways underlying this association **Figure 4D**’s kinase-dimension was transformed into KEGG-pathway space (Kanehisa et al., 2017) (see Supporting Information: Method Details for Figures), yielding a matrix of associations between kinase-signaling pathways and downstream transcriptional programs (**Figure 4E**). This analysis revealed a distinct pattern of transcriptional signatures that distinguished tyrosine-kinase inhibitors, from cell-cycle inhibitors and TGFβ inhibitors (**Figure 4E**). This unsupervised analysis is consistent with the current knowledge of the hierarchical structure of these signaling pathways where RTK-inhibition suppresses MYC-programs and the cell-cycle, while CDK-inhibitors suppress cell-cycle transcriptional programs but not MYC (**Figure 4F**) (Kolch et al., 2015).

### Methodological Summary

Overall, the methods submitted to the final validation round could be broken into three general categories:

1. **Methods relying on a weighted average of differential gene expression and Area Under the Curve (AUC)-based DRC-similarity** across drugs and drug-targets. These included Netphar, SBNB, Xielab, Theragen.
2. **Methods relying on neural networks trained on prior information** relating differential gene expression to drug-targets. These included Atom, DMIS_PDA, and TeamAxolotl
3. **Methods based on fully unsupervised data transformation** combining differential gene expression and DRC data. These included AMbeRland, SenthamizhamV, Signal.

Generally, similarity-weighted average methodologies performed best in SC2 (Netphar 1^st^,SBNB 2^nd^,Theragen 3^rd^)—i.e. they were better at predicting the entire range of targets—while Neural Network-based methodologies performed best in SC1 (Atom 1^st^, DMIS_PDA 3^rd^)—i.e., they were better at predicting targets in the range that could lead to realistic experimental validation. Fully unsupervised methods showed the worst performance. Nonetheless, they still managed to achieve statistical significance, without leveraging any prior knowledge, suggesting the potential for novel mechanistic insight that could be combined with prior knowledge in the future.

In addition, there were differences in the training data sets used by algorithms in the first two categories. While weighted similarity methods used both transcriptional and cytotoxicity data (**Figure 1E & 5A**), neural network methods were trained exclusively with transcriptional profile data (see Sensitivity Data section below). Intriguingly, the winning neural network method (Atom) used protein structure data to further train their neural network (**Figure 5B**). This particular prior knowledge, is worth noting because it underlies several traditional approaches to structure-based drug design (e.g. ligand-docking to homology models) and off-target discovery (e.g. BLAST searches in DrugBank (Wishart et al., 2018)). Unfortunately, while such an approach may eventually help distinguish high-affinity binding targets from key downstream effectors, use of protein sequence information improved Atom’s performance only by a small, non-statistically significant amount.

**Figure 5.**
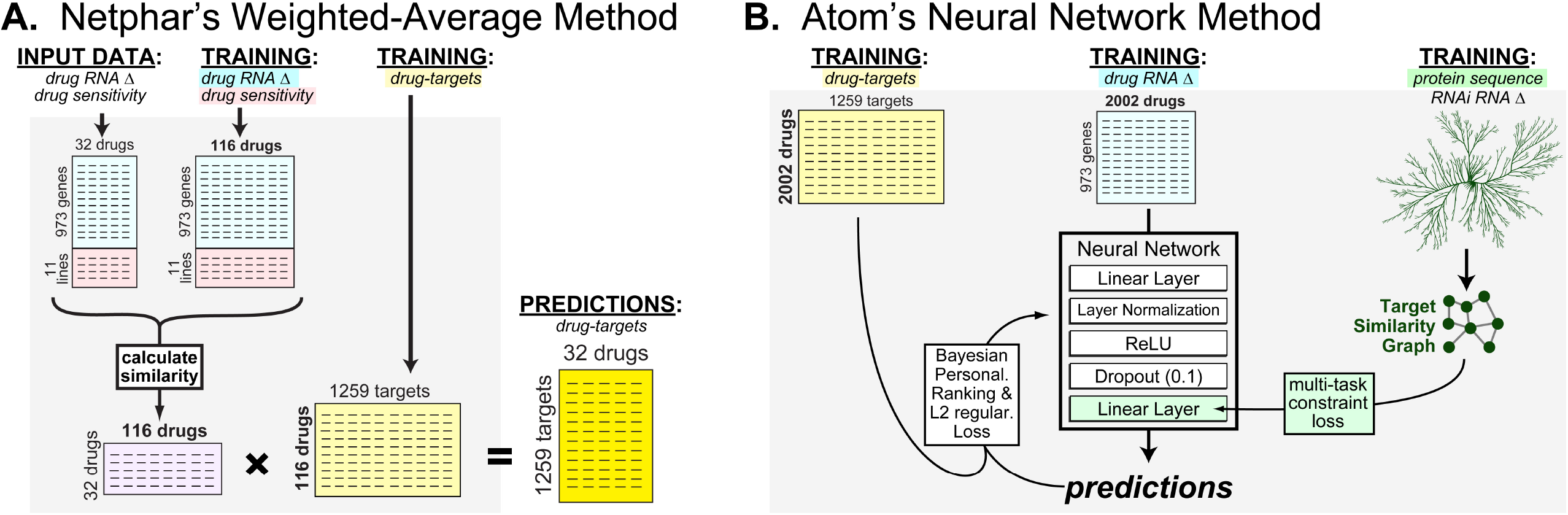
Comparison of the two winning strategies: weighted similarity and neural networks. **A.** Team Netphar (who won SC2) used a simple matrix manipulation procedure to predict drug-targets **B.** Team Atom (who won SC1) used a protein-sequence trained neural network.

### Contribution of Drug Sensitivity Data

Previous work (Szalai et al., 2019) has shown that training on drug-sensitivity profile data can provide comparable prediction performance to training on transcriptional signatures. As such, we sought to investigate the contributions of drug sensitivity and drug transcriptional data to the performance of the winning Netphar model (which utilized both). Drug sensitivity training data was obtained from DrugComb, a curated database that includes batch-corrected drug sensitivities for both single drugs and drug combinations (Zagidullin et al., 2019). In addition to the commonly used IC50, DrugComb provides an AUC-based RI (Relative Inhibition) metric (Malyutina et al., 2019) which captures both the potency and efficacy of drug responses (**Figure S4A**).

Examining correlations between predicted and gold-standard targets, we found that adding drug sensitivity data significantly improved prediction accuracy, relative to transcriptional data alone (**Figure S4b**). In particular, performance improvements were driven by several individual drugs whose targets were poorly predicted based on perturbational profile data only, including sunitinib, crizotinib, and crenolanib (**Figure S4c**). Finally, we tested whether the additional efficacy information provided by the RI metric improved model performance. Indeed, use of the RI metric in the predictive algorithms produced statistically significant, albeit marginal overall improvement (median 0.18 compared to 0.19, paired Wilcoxon test P-value = 0.025), highlighting the potential value of this metric in modeling drug properties.

## DISCUSSION

Mechanism of action elucidation is a critical, yet time-consuming step in the drug development process (Scannell et al., 2012). It helps to identify on- and off-target effects supporting the activity of the compound (polypharmacology), as well as off-target effects that may cause unwanted toxicity. This addresses the two major reasons for clinical trial failure, i.e., lack of safety and/or lack of efficacy (Kola and Landis, 2004; Wehling, 2009). It is reasonable to expect that failure rates may be substantially reduced if compound MoA could be assessed more accurately and comprehensively, including in terms of its tissue-specific differences.

Drug MoA is defined as the set of biochemical interactors and effectors through which the drug produces its pharmacological effects, both positive and negative. These are almost invariably cell-context specific. Despite its relevance, MoA characterization still represents a significant challenge, which is only partially addressed by experimental and computational strategies. Most of the experimental approaches rely on direct binding assays, such as ATP competitive pulldown (Klaeger et al., 2017), affinity purification (Hirota et al., 2012; Ito et al., 2010) or affinity chromatography assays (Aebersold and Mann, 2003). These labor-intensive methods are generally limited to the identification of high-affinity binding targets, rather than the full protein repertoire responsible for compound activity in a tissue, and are often restricted to a specific protein family, such as protein kinases, for instance. Thus critically relevant targets outside of these relatively narrow confines may be missed, as shown by the recent reclassification of the MET tyrosine receptor kinase inhibitor tivantinib as a microtubule inhibitor (Basilico et al., 2013). Indeed, drug polypharmacology is emerging as a critical concept that increasingly impacts our understanding of how drugs work in disease, for instance via a field-effect mediated by multiple targets rather than by their primary, high-affinity binding target. OTS964, for instance, a compound originally developed as a MELK inhibitor, was recently shown to manifest its antitumoral activity via an entirely different target, CDK11, which had originally been missed in its MoA characterization (Lin et al., 2019).

A few computational approaches have also been developed to infer MoA (Keiser et al., 2009; Lomenick et al., 2009; Miller et al., 2002), including using structural and/or genomic information (Yamanishi et al., 2008), text-mining algorithms (Li et al., 2009), or data-mining (Hansen et al., 2009; Perlman et al., 2011). As such, they rely on detailed three-dimensional structures of both the drug molecules and the target proteins or on prior knowledge (literature or database derived) of related compounds. More recently, systematic gene expression profiling (GEP) following compound perturbations in cell lines (Bansal et al., 2014; Lamb et al., 2006; Subramanian et al., 2017; Woo et al., 2015) has furthered the development of computational methods for MoA analysis.

In this manuscript we report on a DREAM community challenge to assess the ability to predict drug mechanism of action inference from drug perturbational profiles, using a comprehensive, experimental protein kinase binding affinity benchmark. This objective benchmark is based on a systematic set of ATP competitive binding assays assessing the ability of 230 candidate kinase inhibitor molecule to bind to one of 255 protein kinases. The corresponding Kinome Binding Repertoire (KBR) database represents the first objective and systematic resource supporting this kind of study.

Drug MoA inference is at best in an embryonic state and the expectation is that this kind of objective assessment through community challenges will significantly increase both the focus on this important topic and provide critical resources for the analysis, potentially leading to a new field of investigation. Consistent with these goals, the results from this study provide unique insight into the approaches that can be used to understand drug MoA, as well as a comprehensive repertoire of datasets and resources that can be leveraged in such studies, as reported by the individual participant labs.

A critical issue emerging from the evaluation of individual prediction performance and individual databases is that the concept of *drug target* is still poorly defined and inconsistent. For instance, even restricting the comparison strictly to protein kinases, comparison of targets defined in DrugBank vs. KBR shows that the former may be missing data and may contain false positive targets, whose binding affinity is >1μM (Figure 3). Yet, it is unclear whether there may be a) false negatives in the KBR, for example if allosteric binding or protein degradation occurred upon drug binding as it would be missed by an ATP competitive binding assay, or b) false positives in DrugBank. More critically, it is unclear whether the targets reported in one database but not the other may actually play a relevant pharmacological role, either in disease treatment or in the emergence of undesirable side effects.

While this was not the main objective of the DREAM Challenge, the study also provides significant insights on the network of effector proteins downstream of high-affinity binding targets. Indeed, the fact that the perturbational signature significantly contributed to correct target inference suggests that the transcriptional regulators that are critical effectors of the high-affinity binding targets represent a valuable reporter assay that can distinguish the MoA of different compounds. Furthermore, the analysis shows that availability of matched DRC and perturbational profile data for each drug provided a statistically significant contribution to the quality of the prediction. Currently, there are databases, such as CTRP (Basu et al., 2013), that provide a DRC profiles for a large number of drugs and cell lines as well as databases such as LINCS (Subramanian et al., 2017) that provide access to large-scale perturbational profile datasets. However, the availability of matched DRC and perturbational profiles, such as those provided by the PANACEA resource, would be beneficial to the community. For instance, drugs such as sunitinib, crizotinib and crenolanib produced significantly poorer performance when analysis was restricted to perturbational profiles but performed significantly better when DRC and perturbational profile data were integrated. These drugs are known to inhibit multiple targets, which is often considered the hallmark of a “dirty drug.” Multiple targets generate a complex profile of downstream effector activation and inactivation, which are harder to deconvolute. To improve these predictions, a set of perturbational data from individual targets, for instance following their CRISPRi-mediated silencing, would be helpful.

Interestingly, tyrosine kinase inhibitors were predicted with higher accuracy by all of the methods (Figure 4A-C). Examining correlations between binding constants and transcriptional profiles, we found that, relative to other inhibitors profiled within the KBR, tyrosine kinases inhibitors were mostly associated with suppression of proliferation signatures (Figure 4D). This is perhaps unsurprising as growth factor control of the cell cycle is typically mediated by receptor tyrosine kinases. Looking at enrichment of KEGG-pathways within Figure 4D’s correlation matrix we were able to identify a decoupling in effects of MYC and the cell-cycle (Figure 4E) that was consistent with hierarchy of known proliferation pathways (Figure 4F). Taken together, these results provide evidence that drug-perturbed transcriptional signatures can retain information on the signaling pathways directly downstream of molecular drug-targets.

While we did not observe major differences between model performance based on modelling strategy, generally, similarity-weighted average methodologies performed best in SC2, while neural networkbased methodologies performed best in SC1. An important insight arising from the challenge is that current methodologies are best at identifying similarities between unknown compounds and compounds already reported in existing databases, rather than at elucidating compound MoA *de novo*. Indeed, all of the methodologies that did not rely on prior databases underperformed when compared to those that did. The fully unsupervised models, without leveraging any prior knowledge, assumed that direct perturbation of a protein leads to the gene expression change of the same protein. While this phenomenon cannot be generally assumed, it led to effective prediction in the case of several targets (e.g.: DRD1, EPHB2 and FYN kinase targeting drugs), suggesting that several proteins can regulate (indirectly) their own expression, probably via a feedback mechanism (Szalai and Saez-Rodriguez, 2020). The fact that all of the proposed methodologies produced statistically significant results suggests that genome wide perturbational profiles bring *de novo* prediction of compound MoA a step closer to be effectively useful in drug discovery.

Overall, this work suggests that predictive models can leverage perturbational data to effectively infer the MoA of small molecules and reveal biological insights about druggable pathways. Future studies using computational modeling to tackle this problem will be critical to the successful application of these methods. Specifically, developing a more systematic knowledge of drug targets, particularly for nonkinase targets, would likely improve the ability of the community to develop accurate models. Additional development and benchmarking of unsupervised prediction methods may also be required for the accurate prediction of targets of novel molecules. Finally, future work will be necessary to elucidate the best practices, limitations, and general applicability of these methods as a step in the drug discovery pipeline.

## ACKNOWLEDGEMENTS

P.A. acknowledges the support of the Generalitat de Catalunya (RIS3CAT Emergents CECH: 001-P-001682 and VEIS: 001-P-001647), the Spanish Ministerio de Economía y Competitividad (BIO2016-77038-R), the European Research Council (SysPharmAD: 614944) and the European Commission (RiPCoN: 101003633). BSz was supported by the Premium Postdoctoral Fellowship Program of the Hungarian Academy of Sciences [460044]. AC was supported by the Cancer Target Discovery and Development Center grant (U01-CA168426) and by NIH shared instrumentation grants S10-OD012351 and S10-OD021764. AC, JG, RA, ED were supported through supplemental funding from the CTD^2^ program (U01-CA217862).

## CONSORTIUM

The members of the CTD^2^ Drug Activity DREAM Challenge Community Consortium are: Renata Retkute, Alidivinas Prusokas, Augustinas Prusokas, Andrea Degasperi, Yasin Memari, João M. L. Dias, Guillermo de Anda-Jáuregui, Santiago Castro-Dau, Cristóbal Fresno, Laura Gómez-Romero, Humberto Gutiérrez-González, Enrique Hernández-Lemus], Soledad Ochoa, José María Zamora-Fuentes, Yue Qiu, Di He, Lei Xie, Gwanghoon Jang, Jungsoo Park, Sungjoon Park, Buru Chang, Sunkyu Kim, Jaewoo Kang, Eugene F. Douglass Jr., Robert Allaway, Bence Szalai, Ron Realubit, Charles Karan, Wenyu Wang, Tingzhong Tian, Adrià Fernández-Torras, Jing Tang, Shuyu Zheng, Alberto Pessia, Ziaurrehman Tanoli, Mohieddin Jafari, Fangping Wan, Shuya Li, Yuanpeng Xiong, Jianyang Zeng, Miquel Duran-Frigola, Martino Bertoni, Pau Badia-i-Mompel, Lídia Mateo, Oriol Guitart-Pla, Patrick Aloy, Verena Chung, Julio Saez-Rodriguez, Justin Guinney, Daniela Gerhard, Andrea Califano (see Supplemental Documentation for consortium author affiliations)

## AUTHOR CONTRIBUTIONS

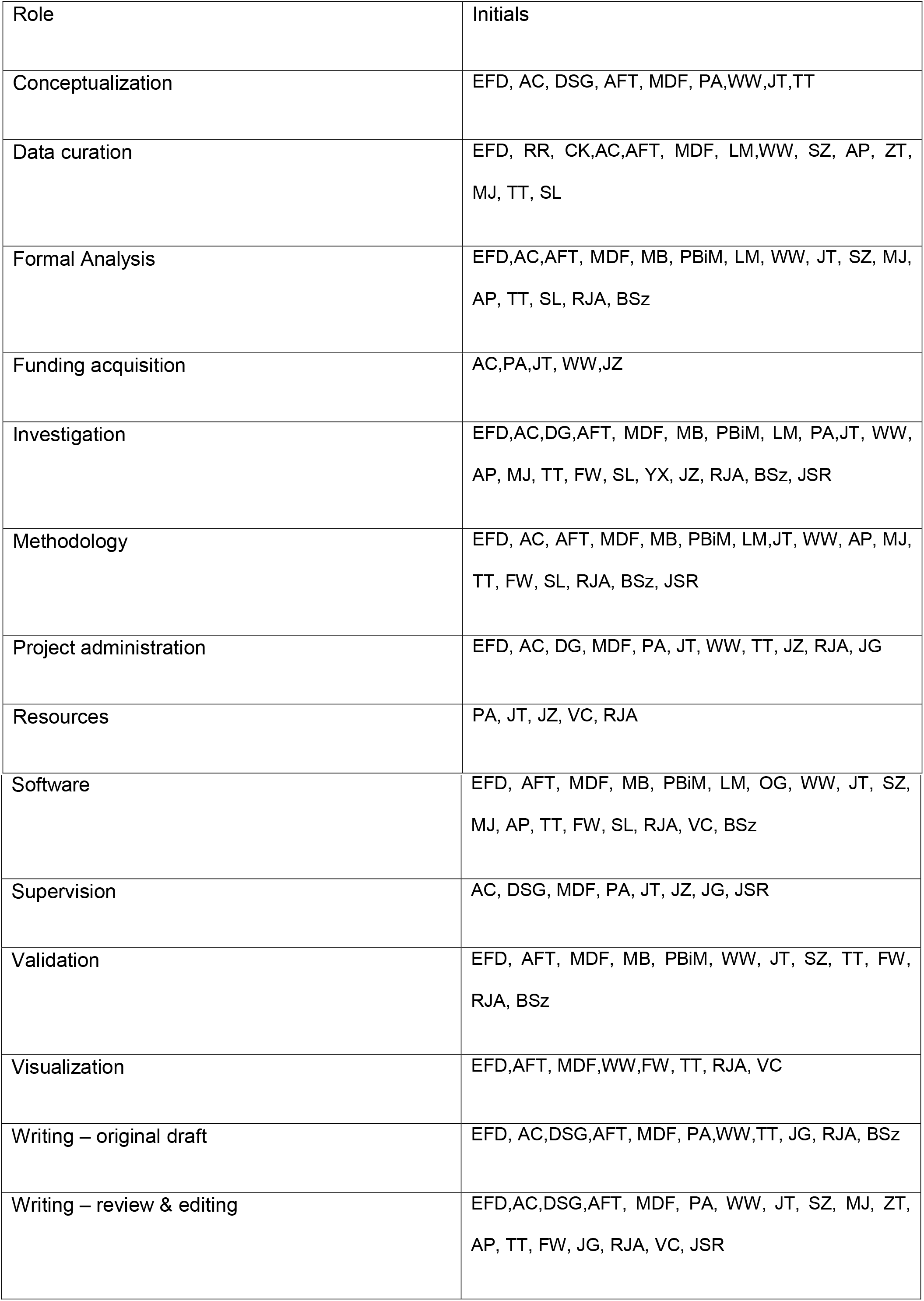

## DECLARATION OF INTERESTS

A.C. is founder, equity holder, and consultant of DarwinHealth Inc., a company that has licensed the PANACEA database used in this manuscript from Columbia University. Columbia University is also an equity holder in DarwinHealth Inc. Other authors declare no conflicts of interest.

## STAR METHODS

### Key Resources Tables

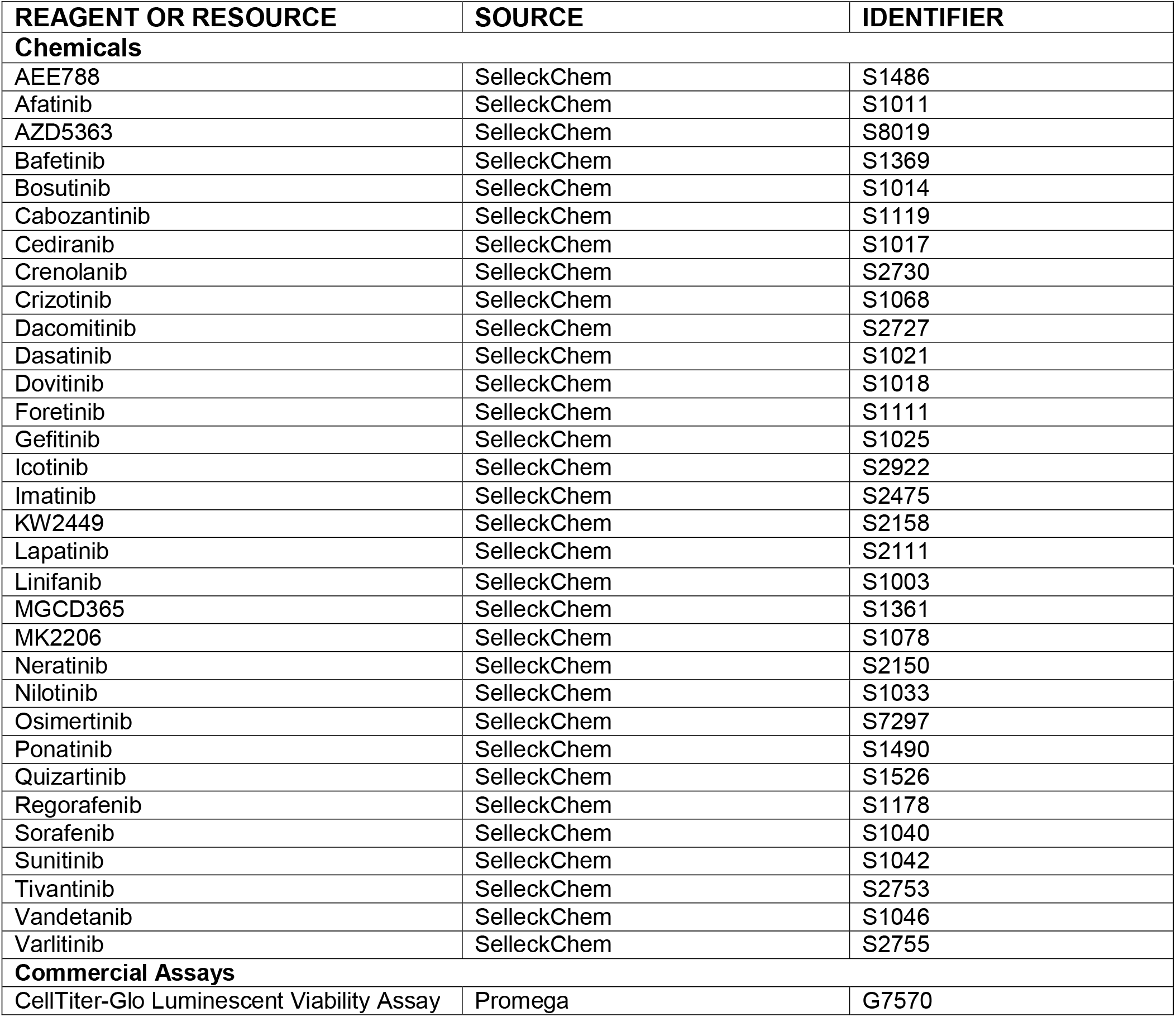

STAR METHODS Key Resources Tables
| REAGENT OR RESOURCE | SOURCE | IDENTIFIER |
| --- | --- | --- |
| **Chemicals** |  |  |
| AEE788 | SelleckChem | S1486 |
| Afatinib | SelleckChem | S1011 |
| AZD5363 | SelleckChem | S8019 |
| Bafetinib | SelleckChem | S1369 |
| Bosutinib | SelleckChem | S1014 |
| Cabozantinib | SelleckChem | S1119 |
| Cediranib | SelleckChem | S1017 |
| Crenolanib | SelleckChem | S2730 |
| Crizotinib | SelleckChem | S1068 |
| Dacomitinib | SelleckChem | S2727 |
| Dasatinib | SelleckChem | S1021 |
| Dovitinib | SelleckChem | S1018 |
| Foretinib | SelleckChem | S1111 |
| Gefitinib | SelleckChem | S1025 |
| Icotinib | SelleckChem | S2922 |
| Imatinib | SelleckChem | S2475 |
| KW2449 | SelleckChem | S2158 |
| Lapatinib | SelleckChem | S2111 |
| Linifanib | SelleckChem | S1003 |
| MGCD365 | SelleckChem | S1361 |
| MK2206 | SelleckChem | S1078 |
| Neratinib | SelleckChem | S2150 |
| Nilotinib | SelleckChem | S1033 |
| Osimertinib | SelleckChem | S7297 |
| Ponatinib | SelleckChem | S1490 |
| Quizartinib | SelleckChem | S1526 |
| Regorafenib | SelleckChem | S1178 |
| Sorafenib | SelleckChem | S1040 |
| Sunitinib | SelleckChem | S1042 |
| Tivantinib | SelleckChem | S2753 |
| Vandetanib | SelleckChem | S1046 |
| Varlitinib | SelleckChem | S2755 |
| **Commercial Assays** |  |  |
| CellTiter-Glo Luminescent Viability Assay | Promega | G7570 |

### Experimental Method Details

The PANACEA database was developed in collaboration between Columbia University Irving Medical Centers (CUIMC)’s High Throughput Screening Center (HTS), Sulzberger Genome Center and the Califano Laboratory in the Department of Systems Biology. Briefly, HTS handled cell-culture, cell-perturbation experiments and RNA extraction; the Genome Center performed RNA sequencing and the Califano laboratory performed data normalization, quality control, benchmarking and scientific and statistical analysis.

#### Cell Line Viability

Cell-lines were obtained from ATCC and cultured using prescribed conditions. To determine optimal seeding density for compound titrations (i.e. cell-growth is linear for the duration of experiment), 3.2 million cells of each cell line were plated and viability measured using CelTiter Glo (Promega Corp.) at 24, 48, 72 and 96 hours. Briefly, 10 mL of 320,000 cells/mL cell-solution was added to column 11 of a 12w deep-well plate. 5mL from column 11 was then serially diluted 1:1 from column 11 through column 2. The Hamilton MicroLab automated liquid handling system’s Cell Line Optimization protocol was used to split the 12 w plates between 4 384 well plates for incubation. 384 well plates were stored in the incubator and at 24, 48, 72 and 96 hours 1 plate was removed and allowed to sit for 15 minutes at room temperature. 25 uL of Cell Titer Glo was added to each well and shaken at 800rpm for 5 min. Finally luminescence was read using the EnVision Multi-Label Reader (Perkin Elmer Inc.). The seeding density which resulted in linear increase of the cells was used for the perturbation experiments.

#### Compound Titration Curves

To determine the 48h ED20 of each drug, cell lines were plated into 96-well tissue culture plates, in 100 μL total volume, and incubated at 37°C. After 16 hours the plates were removed from the incubator and compounds were transferred into assay wells (1 μL) in triplicate. Plates were then returned to the incubator. After 48 hours the assay plates were removed from the incubator and allowed to cool to room temperature prior to the addition of 100 μL of CellTiter-Glo (Promega Inc.) per well. The plates were then mechanically shaken for 5 minutes prior to readout on the EnVision Multi-Label Reader (Perkin Elmer Inc.) using the enhanced luminescence module. Relative cell viability was computed using matched DMSO control wells as reference. ED20 was estimated by fitting a four-parameter sigmoid model to the titration results.

#### Perturbational Profile Generation

Using the previously described plating and perturbation procedure we perturbed each cell-line with each drug at its 48h ED20 value (measured above) or its CMax concentration. In order to optimize the clinical translation potential of the perturbation databases, we used the CMax, defined as the maximum plasma concentration after the administration of the drug at the maximum tolerated dose in patients, (whenever available from published pharmacokinetic studies), as an upper bound for the perturbation studies (**Supp Table 1**). The mRNA from these cells was isolated and profiled by PLATESeq (Nat. Commun. 2017, 8, 105) at 24h after each perturbation.

### Computational Method Details

#### Profile Normalization

RNASeq reads were mapped for each well to the human reference genome assembly 38 using the STAR aligner. Individual plates counts files were then combined, normalized and corrected for batch effects. First, individual counts files were combined across genes and ERCC2 spike-in counts removed, yielding the raw counts file for each cell-line experiment. Second, raw counts were quantile normalized and variance stabilized based on the negative binomial distribution with the DESeq R system package (Love et al., 2014). To account for plate-based batch effects (which are common with drug-perturbed transcriptomic data) normalized expression was batch corrected using ComBat (Johnson et al., 2007).

#### Kinome and PANACEA Data Formatting

Kinome-binding data from Klaeger et. al (Science 2017) was downloaded at https://www.proteomicsdb.org/#projects/4257 via “Supplementary Table 3 Drug Matrices.” Raw data was transformed to −log10 scale and NA’s replaced with the matrix maximum −log10(Kd) of ~4.3 to represent the limit of detection of the technology. PANACEA differential gene expression data was calculated using a moderated Student’s t-test as implemented in the limma package from Bioconductor with respect to pooled DMSO controls across all cell-line plates.

##### Baseline Model

For the baseline model, we used drug perturbation gene expression data from the LINCS-L1000 project (Subramanian et al., 2017) and drug-target information from the Drug Repurposing Hub (Corsello et al., 2017). We calculated consensus signatures (Szalai et al., 2019) for each drug with known target molecules. The DREAM-PanACEA gene expression dataset was standardised using the control measurements, and average signature was calculated for each DREAM-PanACEA drug. We calculated the similarity (Spearman’s correlation) matrix between the LINCS and DREAM-PanACEA drug signatures, using only the measured (landmark) genes of LINCS-L1000. For each DREAM-PanACEA drug, we performed target enrichment (*viper* R package, (Alvarez et al., 2016)) using the drug similarity vector and the known targets of the LINCS drugs. The normalised enrichment scores from target enrichment were further rank normalised for each drug, and submitted as baseline prediction.

##### Scoring Algorithms

Participants submitted predictions for a list of 1259 “druggable” targets and 30 drugs, with each prediction being a confidence score between 0 and 1 (where one is most confident that the target is a true target of a drug). We then filtered each submission to only consider the 255 targets in the gold standard dataset. For the purposes of calculating p-values, we created 1000 null models by generating 1000 random prediction sets. These random predictions were generated by sampling (without replacement) the full set of 1259 “druggable” targets using the dplyr “sample_frac” function to obtain a randomly-ranked set of targets (this procedure was repeated 1000 times) (Wickham et al., 2019). For SC1, we scored teams by evaluating the enrichment of their top 10 predictions for each drug in the gold standard dataset, as well as for one null model prediction, performing a paired Wilcoxon rank sum test (Mann-Whitney test) to generate a p-value for each prediction. We repeated this 10,000 times for each null model to generate a distribution of p-values for each submission, and calculated the mean p-value as the participants’ score. For SC2, the methodology and null models were identical, but instead of evaluating the enrichment of the top 10 predicted targets in the gold standard dataset, we assessed the ranks of the true targets within the full vector of 255 predicted targets for each drug. We again performed a paired Wilcoxon rank sum test (Mann-Whitney test) to generate a p-value for each submission. We repeated this 1000 times for each null model to generate a distribution of p-values for each submission, and calculated the mean p-value as the participants’ score.

Winners were determined by calculating a Bayes factor relative to the top-ranked submission in each category. We bootstrapped all of the submissions that qualified for final scoring by performing 10000 iterations of sampling with replacement for each submission. For each bootstrap, we calculated the p-values as described above to generate a distribution of scores for each submission. Using this distribution of p-values, Bayes factors were calculated for each submission relative to the top-scoring team using the challengescoring R package (https://github.com/sage-bionetworks/challengescoring). Ties were defined as submissions with a Bayes factor <= 3 relative to the top submission.

### Resource Availability

#### Lead Contact

Further information and requests for resources should be directed to and will be fulfilled by the Lead Contact, Andrea Califano.

#### Materials Availability

This study did not generate new unique reagents.

#### Data and Code Availability

Data used in the challenge, submission writeups, and other Challenge resources can be found at www.doi.org/10.7303/syn20968331. Code for scoring the predictions and for generating the null models is available here: https://github.com/Sage-Bionetworks-Challenges/CTD2-Panacea-Challenge, and a Docker container that was used to deploy the scoring algorithm in this challenge is available to all registered Synapse users via the Synapse Docker registry (docker.synapse.org/syn20968331/scoring_harness:v3). A Docker container for team netphar’s model is available at docker.synapse.org/syn21562777/ctd2_netphar_final:final_upd_14March. A Docker container for team Atom’s model is available at docker.synapse.org/syn21560898/ctd2_atom:v1. A Docker container for team SBNB’s model is available at docker.synapse.org/syn21553207/sbnb:9698788.

## Notes

https://www.doi.org/10.7303/syn20968331

## REFERENCES

Aebersold, R., and Mann, M. (2003). Mass spectrometry-based proteomics. Nature 422, 198–207.

Alvarez, M.J., Shen, Y., Giorgi, F.M., Lachmann, A., Ding, B.B., Ye, B.H., and Califano, A. (2016). Functional characterization of somatic mutations in cancer using network-based inference of protein activity. Nat. Genet. 48, 838–847.

Alvarez, M.J., Subramaniam, P.S., Tang, L.H., Grunn, A., Aburi, M., Rieckhof, G., Komissarova, E.V., Hagan, E.A., Bodei, L., Clemons, P.A., et al. (2018). A precision oncology approach to the pharmacological targeting of mechanistic dependencies in neuroendocrine tumors. Nat. Genet.

Anderson, A.C. (2003). The process of structure-based drug design. Chem. Biol. 10, 787–797.

Bansal, M., Yang, J., Karan, C., Menden, M.P., Costello, J.C., Tang, H., Xiao, G., Li, Y., Allen, J., Zhong, R., et al. (2014). NCI-DREAM Community, Gallahan D, Singer D, Saez-Rodriguez J, Xie Y, Stolovitzky G, Califano A, NCI-DREAM Community. A community computational challenge to predict the activity of pairs of compounds. Nat. Biotechnol. 32, 1213–1222.

Barretina, J., Caponigro, G., Stransky, N., Venkatesan, K., Margolin, A.A., Kim, S., Wilson, C.J., Lehár, J., Kryukov, G.V., Sonkin, D., et al. (2012). The Cancer Cell Line Encyclopedia enables predictive modelling of anticancer drug sensitivity. Nature 483, 603–607.

Basilico, C., Pennacchietti, S., Vigna, E., Chiriaco, C., Arena, S., Bardelli, A., Valdembri, D., Serini, G., and Michieli, P. (2013). Tivantinib (ARQ197) displays cytotoxic activity that is independent of its ability to bind MET. Clin. Cancer Res. 19, 2381–2392.

Basu, A., Bodycombe, N.E., Cheah, J.H., Price, E.V., Liu, K., Schaefer, G.I., Ebright, R.Y., Stewart, M.L., Ito, D., Wang, S., et al. (2013). An interactive resource to identify cancer genetic and lineage dependencies targeted by small molecules. Cell 154, 1151–1161.

Bedard, P.L., Hyman, D.M., Davids, M.S., and Siu, L.L. (2020). Small molecules, big impact: 20 years of targeted therapy in oncology. Lancet 395, 1078–1088.

Blumer, K.J., and Johnson, G.L. (1994). Diversity in function and regulation of MAP kinase pathways. Trends Biochem. Sci. 19, 236–240.

Cichonska, A., Ravikumar, B., Allaway, R.J., Park, S., Wan, F., Isayev, O., Li, S., Mason, M., Lamb, A., Tanoli, Z.-U.-R., et al. (2020). Crowdsourced mapping of unexplored target space of kinase inhibitors.

Corsello, S.M., Bittker, J.A., Liu, Z., Gould, J., McCarren, P., Hirschman, J.E., Johnston, S.E., Vrcic, A., Wong, B., Khan, M., et al. (2017). The Drug Repurposing Hub: a next-generation drug library and information resource. Nat. Med. 23, 405–408.

Costello, J.C., Heiser, L.M., Georgii, E., Gönen, M., Menden, M.P., Wang, N.J., Bansal, M., Ammad-ud-din, M., Hintsanen, P., Khan, S.A., et al. (2014). A community effort to assess and improve drug sensitivity prediction algorithms. Nat. Biotechnol. 32, 1202–1212.

Dar, A.C., Das, T.K., Shokat, K.M., and Cagan, R.L. (2012). Chemical genetic discovery of targets and anti-targets for cancer polypharmacology. Nature 486, 80–84.

Duran-Frigola, M., Pauls, E., Guitart-Pla, O., Bertoni, M., Alcalde, V., Amat, D., Juan-Blanco, T., and Aloy, P. (2020). Extending the small-molecule similarity principle to all levels of biology with the Chemical Checker. Nat. Biotechnol.

Günther, S., Kuhn, M., Dunkel, M., Campillos, M., Senger, C., Petsalaki, E., Ahmed, J., Urdiales, E.G., Gewiess, A., Jensen, L.J., et al. (2008). SuperTarget and Matador: resources for exploring drug-target relationships. Nucleic Acids Res. 36, D919–22.

Hansen, N.T., Brunak, S., and Altman, R.B. (2009). Generating genome-scale candidate gene lists for pharmacogenomics. Clin. Pharmacol. Ther. 86, 183–189.

Haverty, P.M., Lin, E., Tan, J., Yu, Y., Lam, B., Lianoglou, S., Neve, R.M., Martin, S., Settleman, J., Yauch, R.L., et al. (2016). Reproducible pharmacogenomic profiling of cancer cell line panels. Nature 533, 333–337.

Hirota, T., Lee, J.W., St John, P.C., Sawa, M., Iwaisako, K., Noguchi, T., Pongsawakul, P.Y., Sonntag, T., Welsh, D.K., Brenner, D.A., et al. (2012). Identification of small molecule activators of cryptochrome. Science 337, 1094–1097.

Hopkins, A.L. (2008). Network pharmacology: the next paradigm in drug discovery. Nat. Chem. Biol. 4, 682–690.

Iorio, F., Bosotti, R., Scacheri, E., Belcastro, V., Mithbaokar, P., Ferriero, R., Murino, L., Tagliaferri, R., Brunetti-Pierri, N., Isacchi, A., et al. (2010). Discovery of drug mode of action and drug repositioning from transcriptional responses. Proc. Natl. Acad. Sci. U. S. A. 107, 14621–14626.

Iorio, F., Knijnenburg, T.A., Vis, D.J., Bignell, G.R., Menden, M.P., Schubert, M., Aben, N., Gonçalves, E., Barthorpe, S., Lightfoot, H., et al. (2016). A Landscape of Pharmacogenomic Interactions in Cancer. Cell 166, 740–754.

Ito, T., Ando, H., Suzuki, T., Ogura, T., Hotta, K., Imamura, Y., Yamaguchi, Y., and Handa, H. (2010). Identification of a primary target of thalidomide teratogenicity. Science 327, 1345–1350.

Johnson, W.E., Li, C., and Rabinovic, A. (2007). Adjusting batch effects in microarray expression data using empirical Bayes methods. Biostatistics 8, 118–127.

Kanehisa, M., Furumichi, M., Tanabe, M., Sato, Y., and Morishima, K. (2017). KEGG: new perspectives on genomes, pathways, diseases and drugs. Nucleic Acids Res. 45, D353–D361.

Kanehisa, M., Sato, Y., Furumichi, M., Morishima, K., and Tanabe, M. (2019). New approach for understanding genome variations in KEGG. Nucleic Acids Res. 47, D590–D595.

Keiser, M.J., Setola, V., Irwin, J.J., Laggner, C., Abbas, A.I., Hufeisen, S.J., Jensen, N.H., Kuijer, M.B., Matos, R.C., Tran, T.B., et al. (2009). Predicting new molecular targets for known drugs. Nature 462, 175–181.

Klaeger, S., Heinzlmeir, S., Wilhelm, M., Polzer, H., Vick, B., Koenig, P.-A., Reinecke, M., Ruprecht, B., Petzoldt, S., Meng, C., et al. (2017). The target landscape of clinical kinase drugs. Science 358.

Kola, I., and Landis, J. (2004). Can the pharmaceutical industry reduce attrition rates? Nat. Rev. Drug Discov. 3, 711–715.

Kolch, W., Halasz, M., Granovskaya, M., and Kholodenko, B.N. (2015). The dynamic control of signal transduction networks in cancer cells. Nat. Rev. Cancer 15, 515–527.

Lamb, J., Crawford, E.D., Peck, D., Modell, J.W., Blat, I.C., Wrobel, M.J., Lerner, J., Brunet, J.-P., Subramanian, A., Ross, K.N., et al. (2006). The Connectivity Map: using gene-expression signatures to connect small molecules, genes, and disease. Science 313, 1929–1935.

Li, J., Zhu, X., and Chen, J.Y. (2009). Building disease-specific drug-protein connectivity maps from molecular interaction networks and PubMed abstracts. PLoS Comput. Biol. 5, e1000450.

Lin, A., Giuliano, C.J., Palladino, A., John, K.M., Abramowicz, C., Yuan, M.L., Sausville, E.L., Lukow, D.A., Liu, L., Chait, A.R., et al. (2019). Off-target toxicity is a common mechanism of action of cancer drugs undergoing clinical trials. Sci. Transl. Med. 11.

Lomenick, B., Hao, R., Jonai, N., Chin, R.M., Aghajan, M., Warburton, S., Wang, J., Wu, R.P., Gomez, F., Loo, J.A., et al. (2009). Target identification using drug affinity responsive target stability (DARTS). Proc. Natl. Acad. Sci. U. S. A. 106, 21984–21989.

Love, M.I., Huber, W., and Anders, S. (2014). Moderated estimation of fold change and dispersion for RNA-seq data with DESeq2. Genome Biol. 15, 550.

Malyutina, A., Majumder, M.M., Wang, W., Pessia, A., Heckman, C.A., and Tang, J. (2019). Drug combination sensitivity scoring facilitates the discovery of synergistic and efficacious drug combinations in cancer. PLoS Comput. Biol. 15, e1006752.

Manning, G., Whyte, D.B., Martinez, R., Hunter, T., and Sudarsanam, S. (2002). The protein kinase complement of the human genome. Science 298, 1912–1934.

Menden, M.P., Wang, D., Mason, M.J., Szalai, B., Bulusu, K.C., Guan, Y., Yu, T., Kang, J., Jeon, M., Wolfinger, R., et al. (2019). Community assessment to advance computational prediction of cancer drug combinations in a pharmacogenomic screen. Nat. Commun. 10, 2674.

Mendez, D., Gaulton, A., Bento, A.P., Chambers, J., De Veij, M., Félix, E., Magariños, M.P., Mosquera, J.F., Mutowo, P., Nowotka, M., et al. (2019). ChEMBL: towards direct deposition of bioassay data. Nucleic Acids Res. 47, D930–D940.

Miller, W.H., Jr, Schipper, H.M., Lee, J.S., Singer, J., and Waxman, S. (2002). Mechanisms of action of arsenic trioxide. Cancer Res. 62, 3893–3903.

Milletti, F., and Vulpetti, A. (2010). Predicting polypharmacology by binding site similarity: from kinases to the protein universe. J. Chem. Inf. Model. 50, 1418–1431.

Perlman, L., Gottlieb, A., Atias, N., Ruppin, E., and Sharan, R. (2011). Combining drug and gene similarity measures for drug-target elucidation. J. Comput. Biol. 18, 133–145.

Proschak, E., Stark, H., and Merk, D. (2019). Polypharmacology by Design: A Medicinal Chemist’s Perspective on Multitargeting Compounds. J. Med. Chem. 62, 420–444.

Saez-Rodriguez, J., Costello, J.C., Friend, S.H., Kellen, M.R., Mangravite, L., Meyer, P., Norman, T., and Stolovitzky, G. (2016). Crowdsourcing biomedical research: leveraging communities as innovation engines. Nat. Rev. Genet. 17, 470–486.

Scannell, J.W., Blanckley, A., Boldon, H., and Warrington, B. (2012). Diagnosing the decline in pharmaceutical R&D efficiency. Nat. Rev. Drug Discov. 11, 191–200.

Schenone, M., Dančík, V., Wagner, B.K., and Clemons, P.A. (2013). Target identification and mechanism of action in chemical biology and drug discovery. Nat. Chem. Biol. 9, 232–240.

Seashore-Ludlow, B., Rees, M.G., Cheah, J.H., Cokol, M., Price, E.V., Coletti, M.E., Jones, V., Bodycombe, N.E., Soule, C.K., Gould, J., et al. (2015). Harnessing Connectivity in a Large-Scale Small-Molecule Sensitivity Dataset. Cancer Discov. 5, 1210–1223.

Shen, Y., Alvarez, M.J., Bisikirska, B., Lachmann, A., Realubit, R., Pampou, S., Coku, J., Karan, C., and Califano, A. (2017). Systematic, network-based characterization of therapeutic target inhibitors. PLoS Comput. Biol. 13, e1005599.

Shoemaker, R.H. (2006). The NCI60 human tumour cell line anticancer drug screen. Nat. Rev. Cancer 6, 813–823.

Subramanian, A., Narayan, R., Corsello, S.M., Peck, D.D., Natoli, T.E., Lu, X., Gould, J., Davis, J.F., Tubelli, A.A., Asiedu, J.K., et al. (2017). A Next Generation Connectivity Map: L1000 Platform and the First 1,000,000 Profiles. Cell 171, 1437–1452.e17.

Szalai, B., and Saez-Rodriguez, J. (2020). Why do pathway methods work better than they should? FEBS Lett.

Szalai, B., Subramanian, V., Holland, C.H., Alföldi, R., Puskás, L.G., and Saez-Rodriguez, J. (2019). Signatures of cell death and proliferation in perturbation transcriptomics data-from confounding factor to effective prediction. Nucleic Acids Res. 47, 10010–10026.

Tang, J., Tanoli, Z.-U.-R., Ravikumar, B., Alam, Z., Rebane, A., Vähä-Koskela, M., Peddinti, G., van Adrichem, A.J., Wakkinen, J., Jaiswal, A., et al. (2018). Drug Target Commons: A Community Effort to Build a Consensus Knowledge Base for Drug-Target Interactions. Cell Chem Biol 25, 224–229.e2.

Wang, Z., Monteiro, C.D., Jagodnik, K.M., Fernandez, N.F., Gundersen, G.W., Rouillard, A.D., Jenkins, S.L., Feldmann, A.S., Hu, K.S., McDermott, M.G., et al. (2016). Extraction and analysis of signatures from the Gene Expression Omnibus by the crowd. Nat. Commun. 7, 12846.

Wehling, M. (2009). Assessing the translatability of drug projects: what needs to be scored to predict success? Nat. Rev. Drug Discov. 8, 541–546.

Wickham, H., Averick, M., Bryan, J., Chang, W., McGowan, L., François, R., Grolemund, G., Hayes, A., Henry, L., Hester, J., et al. (2019). Welcome to the tidyverse. J. Open Source Softw. 4, 1686.

Wishart, D.S., Feunang, Y.D., Guo, A.C., Lo, E.J., Marcu, A., Grant, J.R., Sajed, T., Johnson, D., Li, C., Sayeeda, Z., et al. (2018). DrugBank 5.0: a major update to the DrugBank database for 2018. Nucleic Acids Res. 46, D1074–D1082.

Woo, J.H., Shimoni, Y., Yang, W.S., Subramaniam, P., Iyer, A., Nicoletti, P., Rodríguez Martínez, M., López, G., Mattioli, M., Realubit, R., et al. (2015). Elucidating Compound Mechanism of Action by Network Perturbation Analysis. Cell 162, 441–451.

Yamanishi, Y., Araki, M., Gutteridge, A., Honda, W., and Kanehisa, M. (2008). Prediction of drug-target interaction networks from the integration of chemical and genomic spaces. Bioinformatics 24, i232–40.

Zagidullin, B., Aldahdooh, J., Zheng, S., Wang, W., Wang, Y., Saad, J., Malyutina, A., Jafari, M., Tanoli, Z., Pessia, A., et al. (2019). DrugComb: an integrative cancer drug combination data portal. Nucleic Acids Res. 47, W43–W51.

