## Supplemental Figures for "A Community Challenge for Pancancer Drug Mechanism of Action Inference from Perturbational Profile Data"

### Figure S1

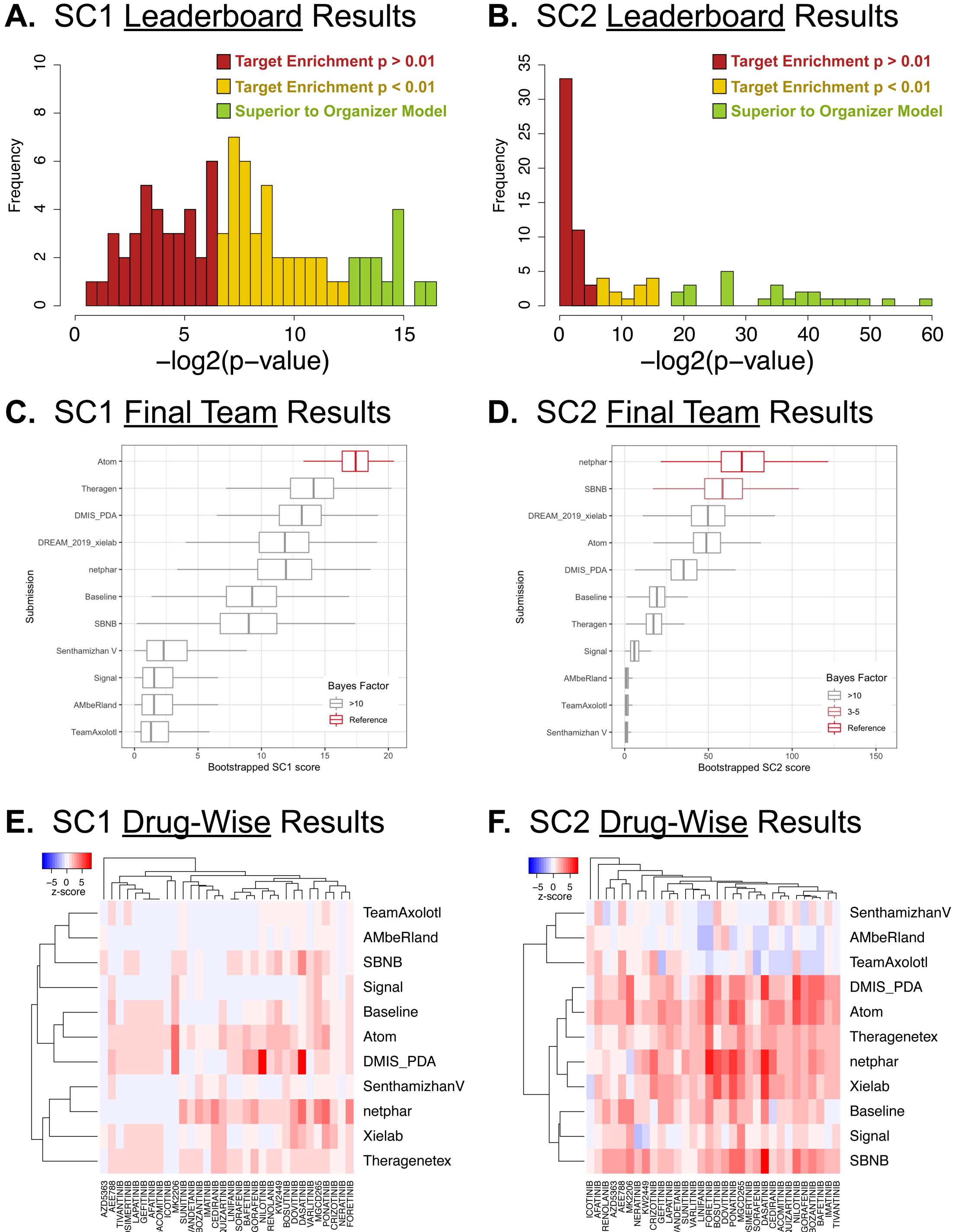
A. SC1 Leaderboard Results
Target Enrichment p > 0.01
Target Enrichment p < 0.01
Superior to Organizer Model
Frequency
−log2(p−value)
B. SC2 Leaderboard Results
Target Enrichment p > 0.01
Target Enrichment p < 0.01
Superior to Organizer Model
Frequency
−log2(p−value)
C. SC1 Final Team Results
Atom
Theragen
DMIS_PDA
DREAM_2019_xielab
netphar
Baseline
SBNB
Senthamizhan V
Signal
AMbeRland
TeamAxolotl
Submission
Bayes Factor
>10
Reference
Bootstrapped SC1 score
D. SC2 Final Team Results
netphar
SBNB
DREAM_2019_xielab
Atom
DMIS_PDA
Baseline
Theragen
Signal
AMbeRland
TeamAxolotl
Senthamizhan V
Submission
Bayes Factor
>10
3-5
Reference
Bootstrapped SC2 score
E. SC1 Drug-Wise Results
z-score
TeamAxolotl
AMbeRland
SBNB
Signal
Baseline
Atom
DMIS_PDA
SenthamizhanV
netphar
Xielab
Theragenetex
F. SC2 Drug-Wise Results
z-score
SenthamizhanV
AMbeRland
TeamAxolotl
DMIS_PDA
Atom
Theragenetex
netphar
Xielab
Baseline
Signal
SBNB

### Figure S2

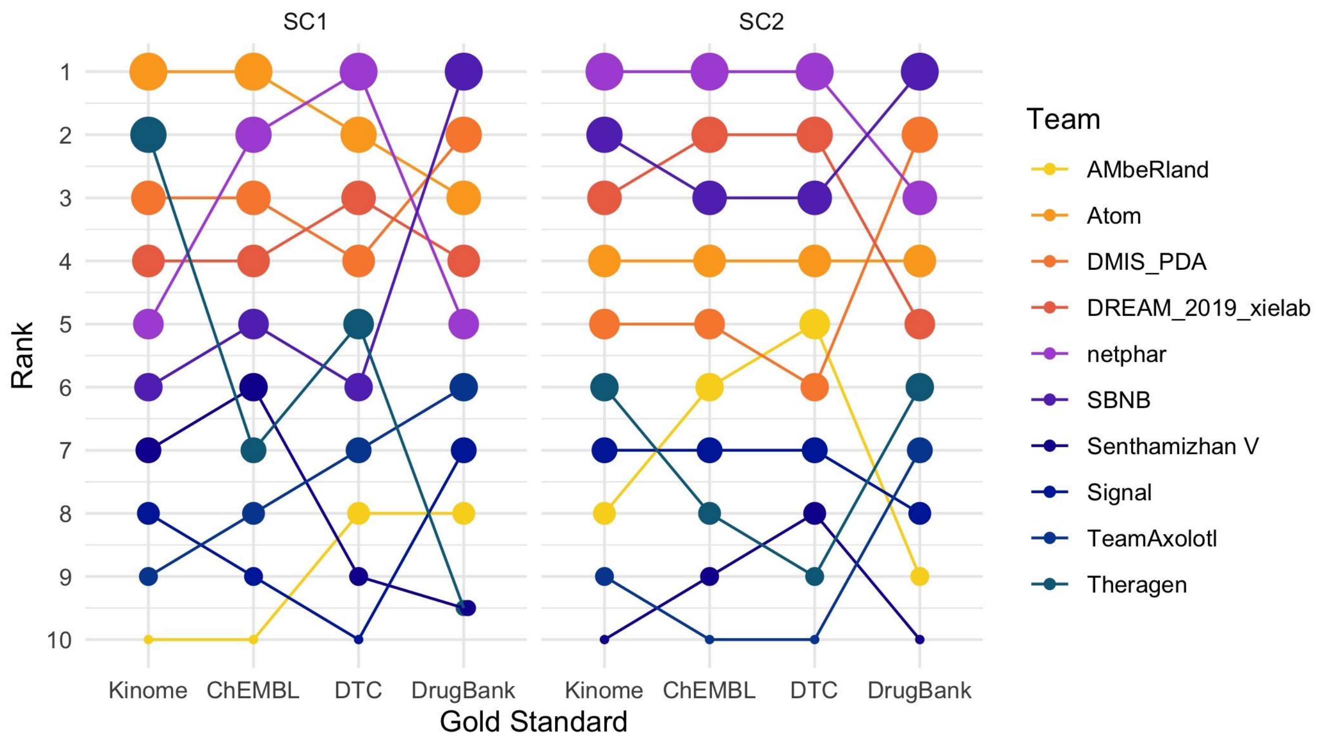
SC1
SC2
Rank
1
2
3
4
5
6
7
8
9
10
Kinome
ChEMBL
DTC
DrugBank
Kinome
ChEMBL
DTC
DrugBank
Gold Standard
Team
AMbeRland
Atom
DMIS_PDA
DREAM_2019_xielab
netphar
SBNB
Senthamizhan V
Signal
TeamAxolotl
Theragen

### Figure S3

#### Average-Scores of **New-Targets** vs **Drug-Bank Targets** in Winning Methods

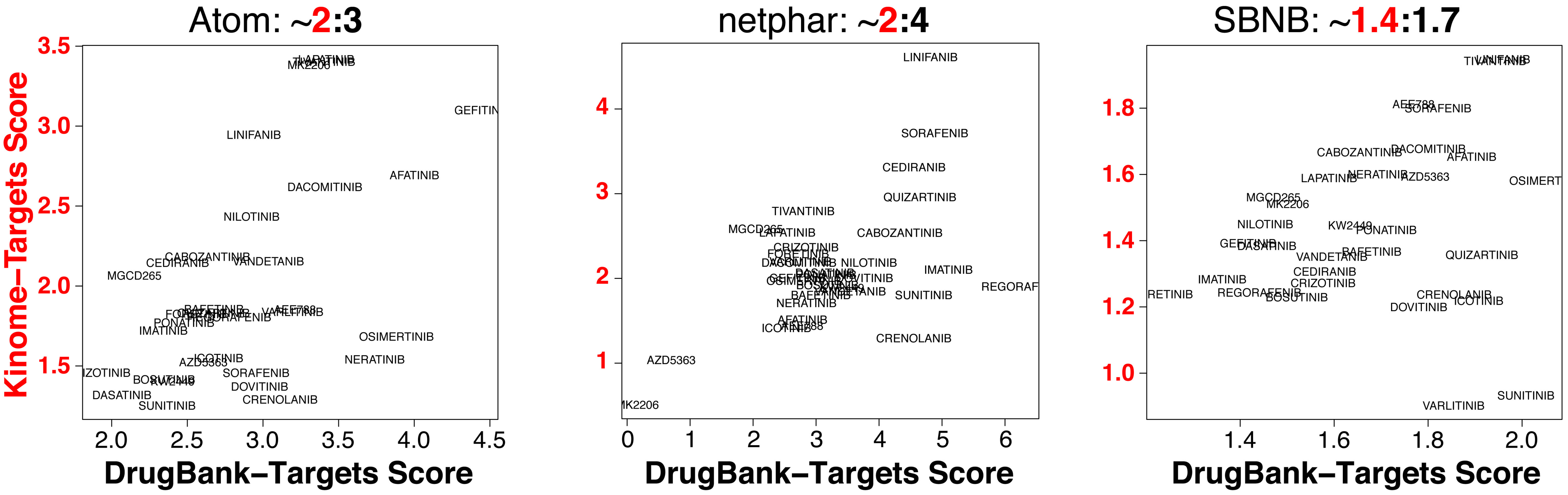

### Figure S4

A.

| Metric | Drug feature | Explaination |
|---|---|---|
| Emax | Efficacy | maximum effect that a drug can produce |
| IC20/IC50 | Potency | amout of a drug needed to produce a given effect |
| AUC | Efficacy + potency | area under the dose response curve |
| RI | | the proportion of AUC to the maximum area or the maximum possible inhibition that one drug can achieve at the same dose range. |

B.

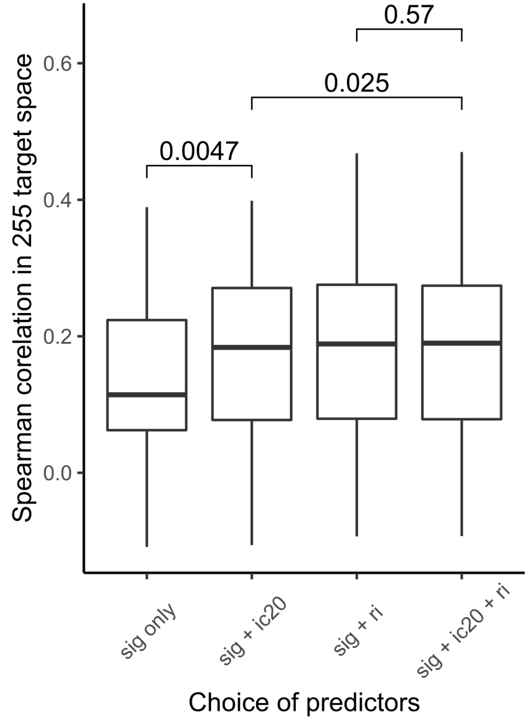


C.

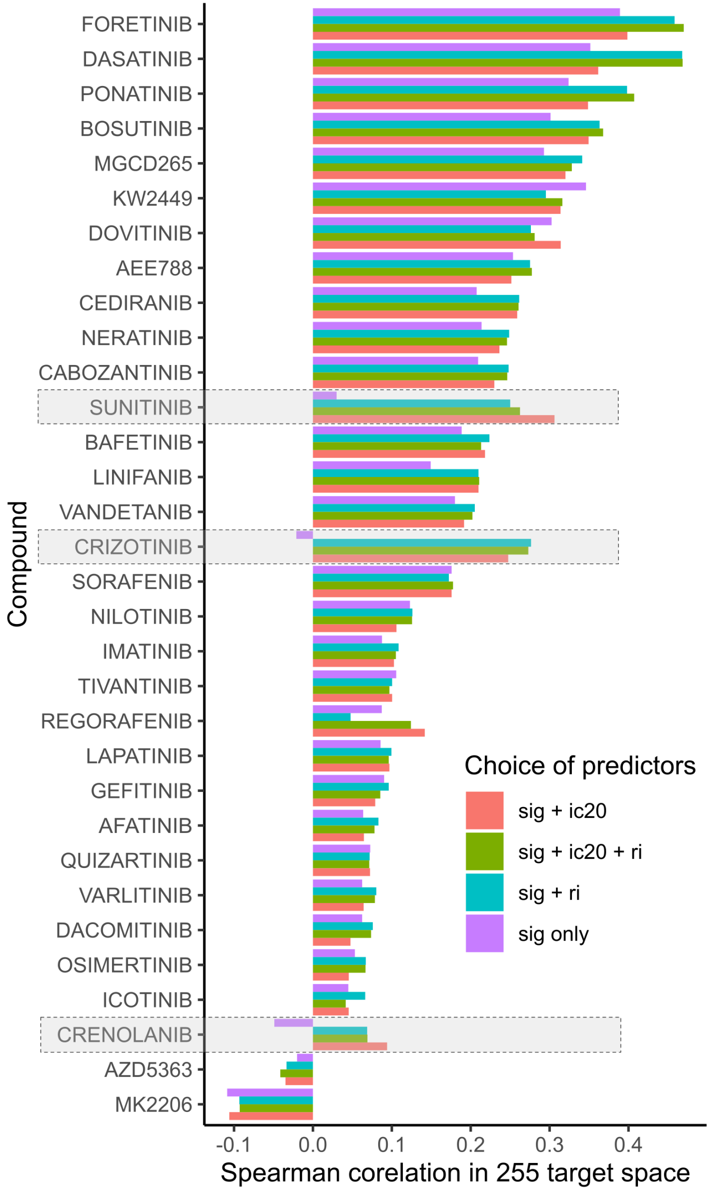

**Figure S5**

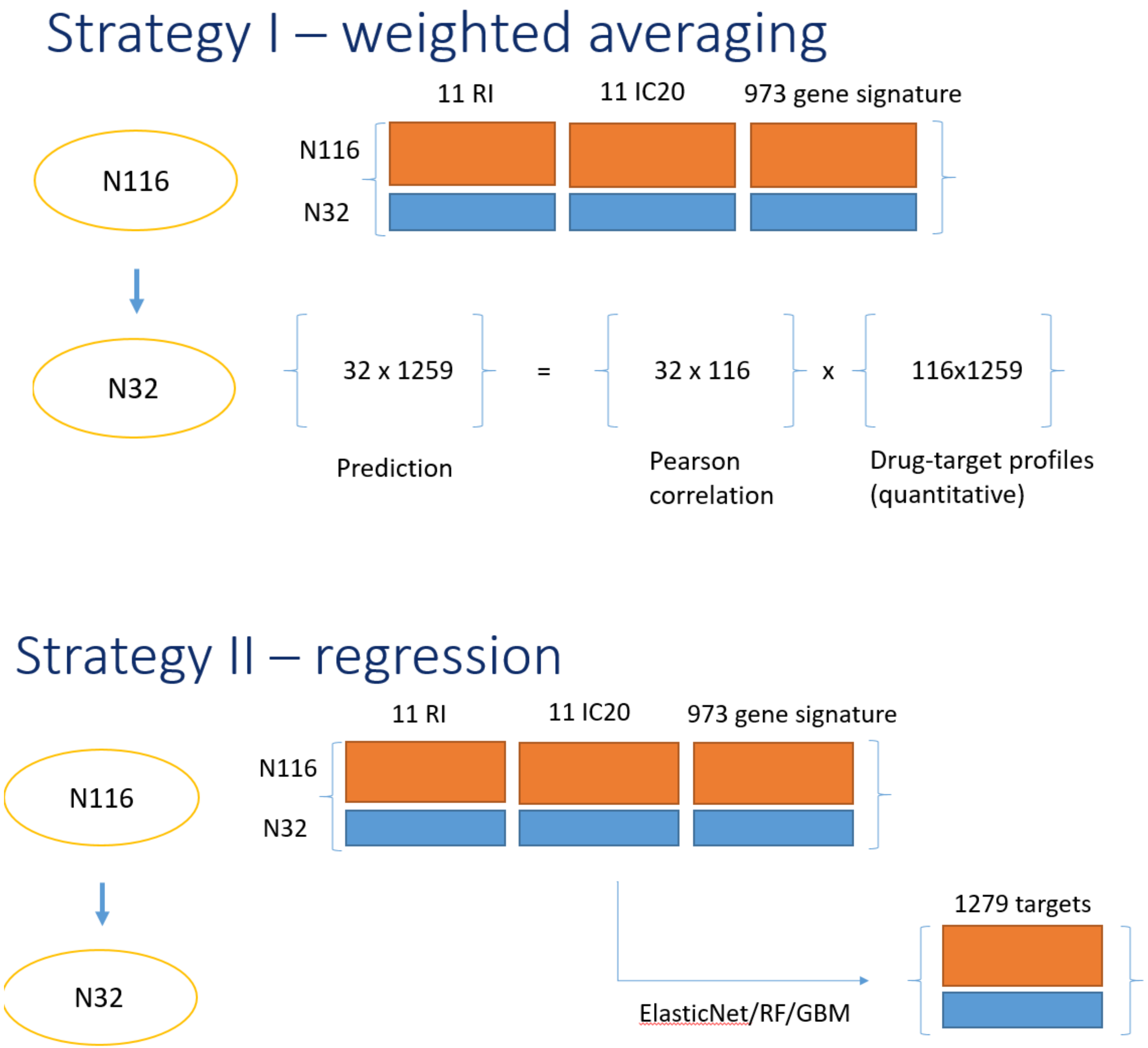
Strategy I – weighted averaging
11 RI
11 IC20
973 gene signature
N116
N116
N32
N32
32 x 1259
=
32 x 116
x
116x1259
Prediction
Pearson
correlation
Drug-target profiles
(quantitative)
Strategy II – regression
11 RI
11 IC20
973 gene signature
N116
N116
N32
N32
1279 targets
ElasticNet/RF/GBM

### Figure S6

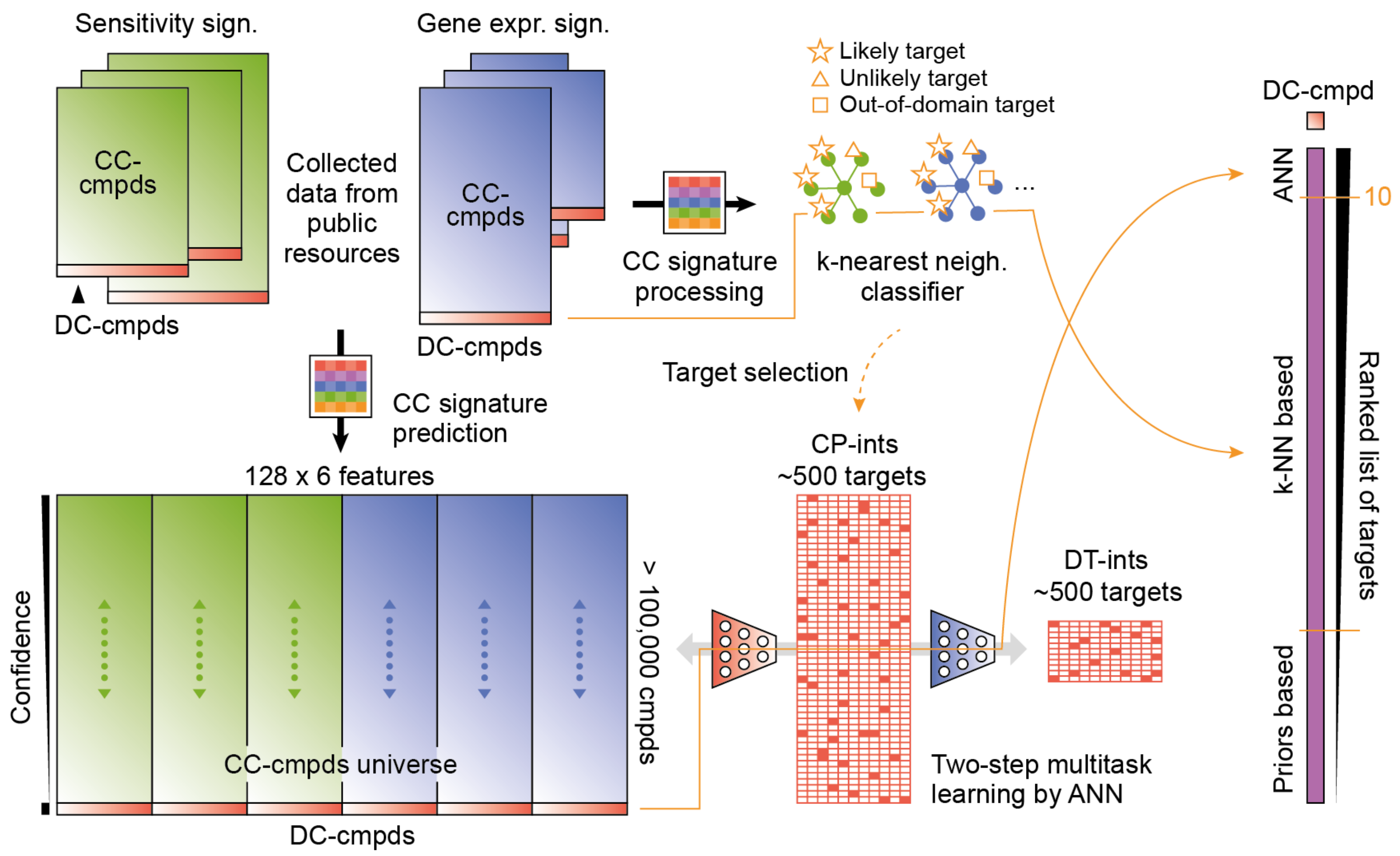

Sensitivity sign.
CC-cmpds
DC-cmpds
Collected data from public resources
Gene expr. sign.
CC-cmpds
DC-cmpds
CC signature processing
Likely target
Unlikely target
Out-of-domain target
k-nearest neigh. classifier
Target selection
CC signature prediction
128 x 6 features
Confidence
CC-cmpds universe
DC-cmpds
> 100,000 cmpds
CP-ints
~500 targets
DT-ints
~500 targets
Two-step multitask learning by ANN
DC-cmpd
ANN
10
k-NN based
Priors based
Ranked list of targets

### Figure S7

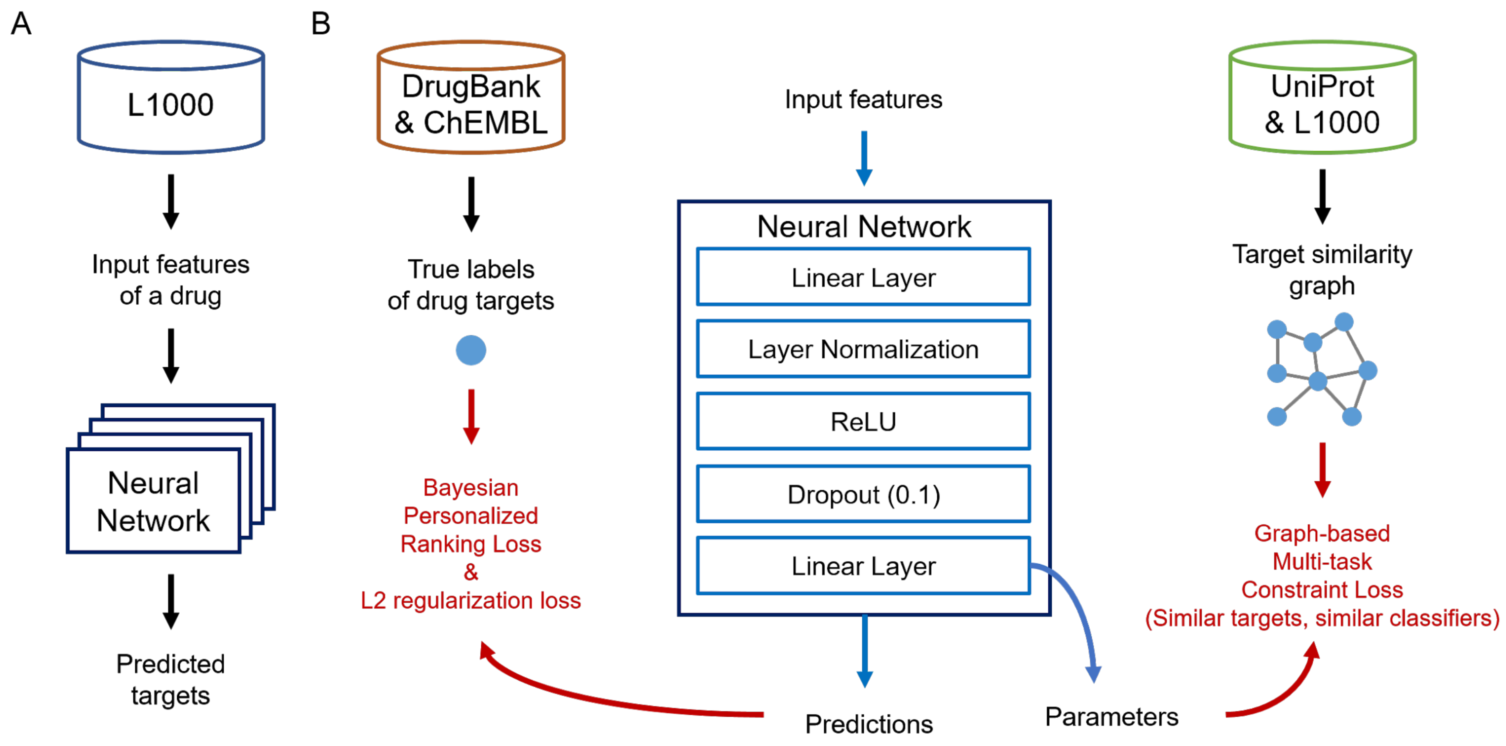

A
L1000
Input features
of a drug
Neural
Network
Predicted
targets
B
DrugBank
& ChEMBL
True labels
of drug targets
Bayesian
Personalized
Ranking Loss
&
L2 regularization loss
Input features
Neural Network
Linear Layer
Layer Normalization
ReLU
Dropout (0.1)
Linear Layer
Predictions
Parameters
UniProt
& L1000
Target similarity
graph
Graph-based
Multi-task
Constraint Loss
(Similar targets, similar classifiers)
