## Supplemental Text for "A Community Challenge for Pancancer Drug Mechanism of Action Inference from Perturbational Profile Data"

‘

### CONSORTIUM AUTHORS AND AFFILIATIONS

These are the members of the CTD[2] Drug Activity DREAM Challenge Community consortium in addition to the byline authors.


**Team AMbeRland: Renata Retkute[1], Alidivinas Prusokas[2], Augustinas Prusokas[3]**
[1] University of Cambridge, Cambrisge, UK
[2] University of Newcastle, Newcastle, UK
[3] Imperial College London, London, UK

**Team Signal: Andrea Degasperi[1,2], Yasin Memari[1,2], João M. L. Dias[1]**
[1] MRC Cancer Unit, University of Cambridge, Hutchison/MRC Research Centre, Cambridge Biomedical Campus, Cambridge, UK,
[2] Academic Laboratory of Medical Genetics, Addenbrooke's Treatment Centre, Addenbrooke's Hospital, Cambridge, UK

**TeamAxolotl: Guillermo de Anda-Jáuregui [1,2,4], Santiago Castro-Dau [1], Cristóbal Fresno [3], Laura Gómez-Romero [1], Humberto Gutiérrez-González [1], Enrique Hernández-Lemus[1,4], Soledad Ochoa [1], José María Zamora-Fuentes [1]**
[1] Computational Genomics Division, National Institute of Genomic Medicine (INMEGEN), México.
[2] National Council on Science and Technology (CONACYT), México.
[3] Technological Development Direction, National Institute of Genomic Medicine (INMEGEN), México.
[4] Computational Complexity Programme, Center for Complexity Sciences at the National Autonomous University of Mexico (C3-UNAM), México.

**Team DREAM_2019_xielab: Yue Qiu[1], Di He[2],Lei Xie[3,4]**
[1] Ph.D. Program in Biology, The Graduate Center, The City University of New York, New York, 10016, United States
[2] Ph.D. Program in Computer Science, The Graduate Center, The City University of New York, New York, NY, 10016, United States
[3] Ph.D. Program in Computer Science, The Graduate Center, and, Department of Computer Science, Hunter College, The City University of New York, New York, NY, 10016, United States
[4] Helen and Robert Appel Alzheimer’s Disease Research Institute, Feil Family Brain & Mind Research Institute, Weill Cornell Medicine, Cornell University, New York, NY, 10065, United States

**Team DMIS_PDA: Gwanghoon Jang [1], Jungsoo Park [2], Sungjoon Park [3], Buru Chang [4], Sunkyu Kim [5], Jaewoo Kang [6]**
[1] Department of Computer Science and Engineering, Korea University, Seoul, Republic of Korea, [2] Interdisciplinary Graduate Program in Bioinformatics, Korea University, Seoul, Republic of Korea

**Team Theragen Bio: Hyunmin Kim [1], Jeein Oh [1], Juyeong Park [1], Seong-Eui Hong [1]**
[1] Theragen Bio Co., Ltd. 4th Fl. Korea Bio Park Bldg. C, 700 Daewangpangyo-ro Bundang-gu, Seongnam-si, Gyeonggi-do Republic of Korea

### SUPPORTING FIGURE LEGENDS:

**Figure S1. Results of community challenge. (A-B)** In the Leaderboard phase, 39 out of 86 models showed statistically significant predictive power by both scoring criteria **(C-D)** In the final round, 6 models beat the base-line model of the organizers **(E-F)** Unsupervised analysis of model performance across individual drugs.

**Figure S2.** Team ranking as a function of gold standard dataset. “Kinome” ranks were determined using the Challenge scoring metrics, gold standard, and null modes generated from the possible kinome targets. ChEMBL, DTC (DrugTargetCommons), and DrugBank ranks were calculated using the DREAM challenge scoring metrics, compound target profiles from each of the respective databases, and using a null model generated from 1259 possible “druggable” targets, to consider targets beyond the kinome. Team ranks are relatively consistent across different gold standards, but in some cases affects the relative ranking of different teams.

**Figure S3.** Comparison method scores for DrugBank-targets (x-axis) versus Unique-Kinome-targets (y-axis) for each drug within the predictions of the winning teams.

**Figure S4 a.** Comparison of different drug sensitivity metrics. **b.** Overall prediction accuracy for the weighted averaging method using different types of predictors. P-values are determined using paired Wilcoxon tests. **c.** Drug-wise prediction accuracy for the weighted averaging method using different predictors. Drughighlighted in grey are those predicted poorly using signature data alone.

**Figure S5.** Overview of Team Netphars modeling Strategy

**Figure S6.** Overview of Team SBNB’s modeling Strategy

**Figure S7.** Model Architecture of Team Atom

### METHOD DETAILS FOR DATA FIGURES IN MAIN TEXT:

**Figure 1 C-D:** PANACEA differential gene expression data was transformed into "Transcriptional Hallmarks" based on definitions of 50 transcriptional signatures defined in (Liberzon et al., 2015). Briefly, an average z-score was calculated for each signature by averaging the z-scores of the individual genes for each signature. PanACEA cell-lines were then averaged to yield a single 32x50 matrix reflecting the relationships of 32 drugs and 50 transcriptional hallmarks. PANACEA and Kinome-binding matrices were then processed for visualization by (1) filtering for the top 30 kinases and signatures by variance and (2) clustering rows and column based on pearson correlation. Filtered data was then visualized using the heatmap.2 function of the gplots package in R. Sidebar annotations of canonical drug-targets were defined based on the DrugBank definitions of drug-targets as detailed in Supporting Table #1.

**Figure 3: (A)** To assess the agreement between DrugBank-Literature and Kinome-binding data, we first defined our reference as: all the kinase-targets defined by DrugBank for our 32-drug library. As the Kinome-binding data gives continuous measurements, it is necessary to define a Kd-threshold to binarize the Kinome-data to compare with DrugBank. For each Kd-threshold, we then calculated the coverage of DrugBank by counting the number of drug-kinase edges identified in the Kinome data and divided by the total number of drug-kinase edges in DrugBank. **(B)** To visualize the new-targets defined in the Kinome-data (but not in DrugBank) we plotted the number of overlapping drug-targets in black (defined in Figure 3A) and newly identified drug-targets in red (defined as NOT being present within DrugBank) for each drug. We then sorted based on total number of targets to aid assessment of polypharmacology.

**Figure 4. (A-B)** To better understand the performance of the three winning models on individual drugs we recalculated team-scored for each drug as z-scores for enrichment (in red) or depletion (in blue) for <uM targets within each drug-vector for both SC1 and SC2. Recalculated scores were sorted by the rank of the average performance across all three teams to identify the drugs which all models performed well on. To better understand the type of inhibitors that models performed the best on we calculated the enrichment of each drugbank target kinases (as defined in Supplmentary Table 1) over the ranked 32-

drug vector in Figures 4A-B using the aREA algorithm in the viper package in R (Alvarez et al., 2016). **(C)** To visualize the kinases sampled by the Klaeger et. al (Klaeger et al., 2017) definitions of drug-targets we color-coded individual kinase-nodes within the Human Kinome phylogenetic tree obtained from the CORAL tool (Metz et al., 2018). Kinases measured in the Kinome-dataset were color coded based the Kinase group that they were a member of as defined in (Manning et al., 2002). **(D)** To better understand the relationships between individual kinases and down-stream transcriptional programs we calculated the correlation matrix between the Kinome Kd's (250 kinases x 84 drugs) and the pan-cancer transcriptional signature PANACEA-data (50 signatures x 84 drugs) across 84 overlapping drugs that occured in both data sets. Correlation matrix was then clustered based on correlation and visualized using the heatmap.2 function in the gplots package in R. Top side-bars were color-coded by kinase groups as defined in (Manning et al., 2002) and colors were chosen to match the Kinome-coverage-phylogenetic tree in Figure 4C. **(E)** To better understand the signalling pathways involved in the kinase-mRNA correlations in Figure 4D, we transformed the individual kinase columns in Figure 4D's correlation matrix into KEGG-defined signalling pathways. This was done by calculating the enrichment of pathway-specific kinases within each transcriptional program vector using the aREA algorithm in the viper package in R. This generated a normalized enrichment score (NES) for each kinase-pathway/mRNA-program pair that is equivalent to a z-score. The visualization on Figure 4E was obtained from the raw pathway-program matrix by slicing top columns associated with receptor tyrosine kinase controlled pathways.

**Figure S3:** To better the relative scores of DrugBank-defined targets and New-Kinome-Data-set defined targets within the winning teams predictions we first normalized all scores by the average-score. The purpose of this was to assure that a random-selection of drug-targets would have a normalized score of 1. For each winning team, and for each drug, we then calculated the average scores of Drug-Bank defined targets and Kinome-defined targets. Encouraging, while DrugBank Targets were consistently higher than Kinome-defined targets, both sets consistently scored better than random sets of drug-targets.

### DETAILED METHODS FOR WINNING TEAM NETPHAR:

**Training Datasets:** The Netphar team collected three types of data related to the compounds: 1) Drug sensitivity data; 2) Drug induced gene signature data and 3) Drug target interaction data. For drug sensitivity data, we utilized the DrugComb database (http://drugcomb.fimm.fi), which is a crowd-sourcing database to collect comprehensive drug sensitivity screen data, including both monotherapy drug screens and drug combination screens (Zagidullin et al., 2019). DrugComb currently consists of drug sensitivity data for 466k combination and 710k monotherapy drug screenings. From DrugComb, we found n = 116 drugs that have dose-response data on at least 7 of the 11 cell lines. Furthermore, for each compound-cell pair, we determined IC20 and RI (relative inhibition, which is based on area under the log10-scaled dose-response curves (Malyutina et al., 2019).) score, as more robust measures for drug sensitivity.

For drug-target interaction data, we utilized the DrugTargetCommons (http://drugtargetcommons.fimm.fi/), which is a crowdsourcing-based database to collectively and manually curate the comprehensive drug-target bioactivity values (Tang et al., 2018). The bioactivity values were transformed into a confidence score between 0 and 1 to indicate the binding affinity potential.

**Analysis of Learning Methods:** To determine the best machine learning models to predict the drug targets, we considered two classes of methods including weighted averaging and regression (Figure below). For weighted averaging, the prediction was made based on the multiplication of the Pearson correlation matrix and the drug-target interaction matrix; while for regression, we considered standard machine learning algorithms including ElasticNet, RandomForest and GBM (Gradient Boosting Machine), for which the model was trained on the n = 116 compounds that were found in DrugComb, and then tested on the n = 32 Challenge compounds. We have utilized the LINCS-L1000 data (Subramanian et al., 2017) to evaluate the methods, and determined the weighted averaging approach that performed better than regression based on 10-fold cross validation.

### DETAILED METHODS FOR WINNING TEAM SBNB:

**Training Data:** As SBNB team, we approached the challenge as a data integration exercise, where we first adapted the transcriptional and sensitivity signatures of the DREAM Challenge compounds to the format of the Chemical Checker (CC) (Duran-Frigola et al., 2020). The CC is a resource that provides processed, harmonized, and ready-to-use bioactivity signatures for about 1M compounds, offering a rich portrait of the small molecule data available in the public domain, and opening an opportunity for making queries that would be otherwise impossible using chemical information alone. The CC expresses bioactivity data as numerical vectors, making them suitable for similarity measurements, clustering, visualization and prediction tasks. Among others, the CC contains cell line sensitivity (Sens) and differential gene expression (DGEx) bioactivity signatures for tens of thousands of compounds (CC compounds), being thus possible to relate this data to the DREAM compounds. To integrate DREAM compounds with CC compounds, we built six different signature types from those bioactivity spaces similar to the ones provided by the DREAM challenge. In three of them, we used growth-inhibition (GI) data of eight cell lines common to the Cancer Therapeutics Response Portal (CTRP, (Seashore-Ludlow et al., 2015)) and the DREAM panel. We then used GI data as features to train a classifier to infer the expected CTRP sensitivity (Sens) profile of DREAM compounds, as well as biomarkers and annotations from the PharmacoDB resource (Smirnov et al., 2018). Thus, we could connect the DREAM compounds to the hundreds of drugs available in the public drug sensitivity panels. Likewise, DREAM DGEx data was integrated with LINCS DGEx (Level 5) signatures (Subramanian et al., 2017), along with the Touchstone reference collection of perturbational profiles. Additionally, we mapped the DREAM and LINCS DGEx to a collection of manually curated expression signatures from Gene Expression Omnibus (CREEDS, (Wang et al., 2016)), in order to capture cell-unspecific profiles, since only one cell line was shared between the DREAM and LINCS L1000 resources. We moved from individual gene expression to global expression signatures with the aim of capturing possible transcriptional regulatory programs shared among the compounds, enabling thus a more comprehensive integration of the DREAM and LINCS datasets.

Once we contextualised DREAM compounds within the larger CC compounds collection, we used the Sens/DiffGEx signatures as input for conventional target prediction methods, based on previously known ligand-binding profiles. In brief, to prevent overfitting due to the limited number of CC-exp compounds, we first used CC signatures to train a k-nearest neighbors (kNN) classifier to identify the most probable targets for each DREAM compound.

**Similarity Analysis:** We simply looked in DrugBank for CC compounds having similar signatures to the DREAM compounds, and suggested the CC annotated targets as putative targets for the DREAM compounds. Then, in a second step, we used a much larger set of over 100k bioactive compounds in the Chemical Checker, for which we inferred their gene expression and cell line sensitivity signatures to train a multitask, quality-aware artificial neural network (ANN) primarily based on chemogenomics data (i.e. compound interactions) from ChEMBL (Gaulton et al., 2017) and refined it with DrugBank drug-target data (Wishart et al., 2018). Finally, the top predictions of this network were used to reorganize the top-10 kNN targets for each drug.

### DETAILED METHODS FOR WINNING TEAM ATOM:

**Input features:** For each compound, its compound-perturbed gene expression feature was calculated from Level 5 data of the LINCS L1000 platform of phase I (GSE92742) and phase II (GSE70138). To obtain a consensus feature for each compound without considering other conditions like cell line, dose and time, all the Level 5 signatures corresponding to the same compound were selected and averaged using MODZ algorithm introduced in L1000 paper (Subramanian et al., 2017). In order to suit for the challenge, we compared the RNA-seq data with the L1000 data and selected 973 overlapping genes as input features.

**Target similarity graph:** During the model training, a graph-based multi-task constraint was used to train our model (described below). The target similarity graph was constructed by using two types of metrics, including a sequence similarity from protein primary sequences as well as a genomic similarity from gene knockdown perturbed gene expression profiles. The protein primary sequences were first obtained from UniProt database according to their gene IDs. Then, the Smith-Waterman sequence alignment scores were computed by an alignment tool (https://github.com/mengyao/Complete-Striped-Smith-Waterman-Library). The sequence similarity between two proteins was then defined as the normalized alignment scores, that is, $\frac{sw(s_1,s_2)}{\sqrt{sw(s_1,s_1)\times sw(s_2,s_2)}}$, where $sw(s_1, s_2)$ stands for the alignment score between protein sequences $s_1$ and $s_2$. The gene knockdown perturbed gene expression profiles were obtained from the L1000 database and processed using the same protocol as drug features described above. The genomic similarity between two targets was defined as $max\{0, r(e_1, e_2)\}$, where $r(e_1, e_2)$ stands for Pearson's correlation coefficient between gene expression profiles $e_1$ and $e_2$. Finally, we averaged these two matrices and constructed a K-nearest neighbor (KNN) (K=10) graph as our final target similarity graph.

As the problem is to predict the potential targets for a compound/drug of interest, we formulate this problem as a multi-label classification problem, where the input of a compound is the compound-perturbed gene expression feature $x \in R^{973}$ derived from LINCS L1000 platform, and the output is a binary vector $y \in R^{769}$ indicating the binding probabilities to 769 pre-defined protein targets. We used an ensemble of neural networks to make predictions.

**Neural Network Architecture:** We use an ensemble of single-layer neural networks to model the relationship between $x$ and $y$. The model architecture of each base learner (i.e., a single-layer neural network) is shown in Figure XX. For each base learner, three losses are used to train its parameters. The first one is Bayesian Personalized Ranking (BPR) loss (Rendle et al., 2012). Specifically, let $S_{i,j}$ denote the predicted score between drug $i$ and protein $j$ produced by our model. Then, BPR loss is defined as:

$BPR\_Loss = - log\ log(S_{i,j} - S_{i,k})$, where protein $j$ is the known target of drug $i$ while protein $k$ is not. During neural network training, we sampled a batch (batch size = 256) of drugs, and for each drug $i$, we sampled pairs $(i, j)$ and $(i,k)$ to perform forward and backward propagation. The second loss is a multi-task constraint loss (Zhou et al., 2011). The multi-task constraint loss is defined as: $Multitask\_Loss = trace(WLW^T)$, where $W$ is the learnable parameter of the last layer of the neural network, $L$ is the normalized graph laplacian of the target similarity graph defined above. This loss encourages the similar targets to have similar classifiers. The last loss is the weight decay (i.e., L2_regularization) for controlling the model complexity. The combined loss is defined as BPR_Loss $+\lambda_1 Multitask\_Loss+$ $\lambda_2$L2_regularization, where $\lambda_1$ and $\lambda_2$ are used to balance different losses. This combined loss was optimized by Adam optimizer with learning rate = 0.001.

We used 10-fold cross validation to train models. For each fold, 1/10 of the drugs were used as test data. Among the remaining 9/10 drugs, 1/10 of drugs were left out as validation data and the rest drugs were used as training data. This strategy was used to perform hyperparameter selection (i.e., dropout rate, $\lambda_1$, $\lambda_2$, hidden size of the neural network and training epoch). During training, early stopping was used to prevent overfitting. For each epoch, we compared the model performance on validation data with the best performance. The training process would be stopped as long as the performance on the validation data no longer improves in consecutive 100 epochs.

We used ensemble learning approach to further boost the performance. We constructed 100 different neural network models from { $\lambda_1$= 0.0001, 0.00001} × { $\lambda_2$= 0.0001} × {hidden size of neural network = 256, 512, 1024, 2048, 4096} × {10 different folds}. These hyperparameter ranges produced decent prediction performance during our hyperparameter selection. We then averaged the prediction scores from these models to produce the final scores.

Supplementary References:

Alvarez, M.J., Shen, Y., Giorgi, F.M., Lachmann, A., Ding, B.B., Ye, B.H., and Califano, A. (2016). Functional characterization of somatic mutations in cancer using network-based inference of protein activity. Nat. Genet. *48*, 838–847.

Duran-Frigola, M., Pauls, E., Guitart-Pla, O., Bertoni, M., Alcalde, V., Amat, D., Juan-Blanco, T., and Aloy, P. (2020). Extending the small-molecule similarity principle to all levels of biology with the Chemical Checker. Nat. Biotechnol.

Gaulton, A., Hersey, A., Nowotka, M., Bento, A.P., Chambers, J., Mendez, D., Mutowo, P., Atkinson, F., Bellis, L.J., Cibrián-Uhalte, E., et al. (2017). The ChEMBL database in 2017. Nucleic Acids Res. *45*, D945–D954.

Klaeger, S., Heinzlmeir, S., Wilhelm, M., Polzer, H., Vick, B., Koenig, P.-A., Reinecke, M., Ruprecht, B., Petzoldt, S., Meng, C., et al. (2017). The target landscape of clinical kinase drugs. Science *358*.

Liberzon, A., Birger, C., Thorvaldsdóttir, H., Ghandi, M., Mesirov, J.P., and Tamayo, P. (2015). The Molecular Signatures Database (MSigDB) hallmark gene set collection. Cell Syst *1*, 417–425.

Malyutina, A., Majumder, M.M., Wang, W., Pessia, A., Heckman, C.A., and Tang, J. (2019). Drug combination sensitivity scoring facilitates the discovery of synergistic and efficacious drug combinations in cancer. PLoS Comput. Biol. *15*, e1006752.

Manning, G., Whyte, D.B., Martinez, R., Hunter, T., and Sudarsanam, S. (2002). The protein kinase complement of the human genome. Science *298*, 1912–1934.

Metz, K.S., Deoudes, E.M., Berginski, M.E., Jimenez-Ruiz, I., Aksoy, B.A., Hammerbacher, J., Gomez, S.M., and Phanstiel, D.H. (2018). Coral: Clear and Customizable Visualization of Human Kinome Data. Cell Syst *7*, 347-350.e1.

Rendle, S., Freudenthaler, C., Gantner, Z., and Schmidt-Thieme, L. (2012). BPR: Bayesian Personalized Ranking from Implicit Feedback.

Seashore-Ludlow, B., Rees, M.G., Cheah, J.H., Cokol, M., Price, E.V., Coletti, M.E., Jones, V., Bodycombe, N.E., Soule, C.K., Gould, J., et al. (2015). Harnessing Connectivity in a Large-Scale Small-Molecule Sensitivity Dataset. Cancer Discov. *5*, 1210–1223.

Smirnov, P., Kofia, V., Maru, A., Freeman, M., Ho, C., El-Hachem, N., Adam, G.-A., Ba-Alawi, W., Safikhani, Z., and Haibe-Kains, B. (2018). PharmacoDB: an integrative database for mining in vitro anticancer drug screening studies. Nucleic Acids Res. *46*, D994–D1002.

Subramanian, A., Narayan, R., Corsello, S.M., Peck, D.D., Natoli, T.E., Lu, X., Gould, J., Davis, J.F., Tubelli, A.A., Asiedu, J.K., et al. (2017). A Next Generation Connectivity Map: L1000 Platform and the First 1,000,000 Profiles. Cell *171*, 1437-1452.e17.

Tang, J., Tanoli, Z.-U.-R., Ravikumar, B., Alam, Z., Rebane, A., Vähä-Koskela, M., Peddinti, G., van Adrichem, A.J., Wakkinen, J., Jaiswal, A., et al. (2018). Drug Target Commons: A Community Effort to Build a Consensus Knowledge Base for Drug-Target Interactions. Cell Chem Biol *25*, 224-229.e2.

Wang, Z., Monteiro, C.D., Jagodnik, K.M., Fernandez, N.F., Gundersen, G.W., Rouillard, A.D., Jenkins, S.L., Feldmann, A.S., Hu, K.S., McDermott, M.G., et al. (2016). Extraction and analysis of signatures from the Gene Expression Omnibus by the crowd. Nat. Commun. *7*, 12846.

Wishart, D.S., Feunang, Y.D., Guo, A.C., Lo, E.J., Marcu, A., Grant, J.R., Sajed, T., Johnson, D., Li, C., Sayeeda, Z., et al. (2018). DrugBank 5.0: a major update to the DrugBank database for 2018. Nucleic Acids Res. *46*, D1074–D1082.

Zagidullin, B., Aldahdooh, J., Zheng, S., Wang, W., Wang, Y., Saad, J., Malyutina, A., Jafari, M., Tanoli, Z., Pessia, A., et al. (2019). DrugComb: an integrative cancer drug combination data portal. Nucleic Acids Res. *47*, W43–W51.

Zhou, Jiayu, Jianhui Chen, and Jieping Ye (2011). Malsar: Multi-task learning via structural regularization. Arizona State University 21 (2011).
